# Metastatic founder cell candidates resemble preimplantation embryonic blastomeres

**DOI:** 10.64898/2026.08.28.747818

**Authors:** Huiqin Körkel-Qu, Elia Raya, Miodrag Guzvic, Christoph Irlbeck, Tobias Mederer, Daniel Spitzl, Zbigniew Czyz, Leon Schunicht, Stephan Seitz, Jonas Roth, Brigitte Rack, Nadia Harbeck, Hasan Kurdieh, Roman Mayr, Max Burger, Tobias Robold, Hans-Stefan Hofmann, Markus Weber, Matthias Maak, Klaus-Peter Janssen, Sarah Hücker, Stefan Kirsch, Melanie Werner-Klein, Anthony C.F Perry, Christoph A. Klein

## Abstract

Disseminated cancer cells (DCC) in non-metastatic carcinoma patient bone marrow (BM) are predictive of metastasis. Those detected by epithelial cytokeratin or EpCAM expression have poorly-characterized transcription profiles due to their extreme rarity: 1∼2 cells per 2x 10^6^ BM cells in every third non-metastatic patient. We here characterize the transcriptomes of DCCs. Single-cell RNA-sequencing (scRNA-seq) of 864 EpCAM-positive cells (from 1,151 cancer patients) in BM or lymph nodes (LN) revealed plasma, immune, myeloid, erythroid progenitor cells and two candidate DCC populations, termed M0-DCC and M1-DCC. M0-DCC, mostly from non-metastatic M0-stage patients, displayed the highest known adult stemness scores, and were transcriptomically reminiscent of human cleavage-stage, preimplantation embryos. M1-DCC represented cancer cells undergoing the epithelial-mesenchymal transition (EMT), corresponding to later, implanting and gastrulating embryos. Detection of early-embryo-like DCC categorised patients at highest risk for metastatic progression. Furthermore, high M0-DCC scores predicted the metastatic potential of human cell lines from the Cancer Cell Line Encyclopedia. M0-DCC gene expression profiles can be reversibly induced from M1-DCC-like cells *in vitro*. The close correspondence between gene expression profiles in immediate early embryonic development and metastatic founder cell candidates provides strong evidence that the onset of cancer and metastasis recruits mechanisms employed in fertilization.

## Introduction

Once metastatic disease becomes detectable by clinical imaging (i.e. typically when tumours are 0.5-1 cm in diameter) it is mostly incurable: recent clinical development such as targeted immunotherapies have resulted in life-span extension, but few cures ^1^. Therefore, much hope rests on detecting single disseminated cancer cells (DCC) in bone marrow (BM) months to years before clinical manifestation ^2^, as DCC promise to provide hints on how to prevent the outgrowth of metastasis. The problem has been that due to their extreme rareness, DCC remain poorly characterized. However, the advent of single-cell technologies has enabled DCC genomic and transcriptomic profiling ^3,4^, leading to the finding that they disseminate very early in tumorigenesis, that they acquire many genetic features of cancer outside of the primary tumour, and that metastatic colony formation is accompanied by significant changes in gene expression related to phenotypic adaptation ^5–10^.

Clinical follow-up studies have repeatedly demonstrated that DCC predict clinical outcomes for a large variety of carcinomas ^11^ and melanoma ^5,12^. In carcinomas, two markers were predominantly used to detect DCC with high specificity: epithelial cytokeratins in mesenchymal organs like BM, or epithelial cell adhesion molecule, EpCAM, in lymph nodes (LN). However, as employed, these markers suffer from shortcomings that include, (i) a substantial portion (depending on marker and cancer type) of DCC-negative patients progress to metastasis; (ii) not all patients with DCC progress, and (iii) the hazard ratio imposed by DCC has been modest and varied between cancers with clear DCC impact, for example in breast cancer, and conflicting data for other carcinomas ^11,13,14^. Altogether, clinical data suggest that typical detection assays miss relevant DCC-subtypes, and that biological processes such as EMT or other manifestations of cellular plasticity blur assessment of early systemic cancer.

To better characterize DCC, we set out to identify DCC in BM from breast, prostate, oesophageal cancer, and in BM and LN from non-small cell lung cancer (NSCLC) patients, using high EpCAM protein expression as the sole selection criterion. Although previous reports had indicated that EpCAM is not exclusive to tumour cells in BM ^4,15^, we chose this marker for several reasons. First, EpCAM is expressed in epithelial organs and most carcinomas derived from them ^16^. Secondly, EpCAM expression is higher in epithelial cells compared to EpCAM-expressing non-epithelial cells ^17^, providing a rationale for distinguishing between the two populations. Thirdly, EpCAM is highly expressed in the cells of preimplantation embryos, indicating that EpCAM reflects a founding state of relatively high cellular potency and that de-differentiation during disease progression is not necessarily linked to EpCAM loss16. We therefore decided to consider each BM or LN EpCAM+ cell as a *bona fide* DCC and accordingly re-classified all cells after in-depth transcriptome profiling by single-cell RNA-sequencing (scRNA-seq). In this way, we sought to separate true DCC from healthy BM cells and thereby access their gene expression profiles.

## Results

### DCC detection using EpCAM

We obtained BM samples from breast cancer (n=472), prostate cancer (n=489), oesophageal cancer (n=70) patients, and BM and LN samples from NSCLC patients (n=120; supplementary table 1). Enrichment for epithelial cells followed by EpCAM staining and screening, yielded 1,388 EpCAM+ cells that were subjected to quality control (QC) and scRNA-seq (Fig 1A and supplementary table 1). Patient samples fell into one of three clinical categories: Union for International Cancer Control (UICC) stage M0 (no evidence of metastasis); UICC stage M1 (clinical manifestation of metastasis); control, age-matched non-cancer patients undergoing trauma or hip replacement surgery. Of all isolated EpCAM+ cells, 864 were subjected non-selectively to further analysis. Since previous analyses and our preliminary data had indicated that EpCAM is expressed in thymic plasma cells ^18^ and erythroid progenitor cells (ERP) ^15^ in BM, we added respective markers, CD27/CD319 or CD235a, to identify double-positive, putative plasma cells or ERP for subsequent profiling annotation.

**Figure 1:**
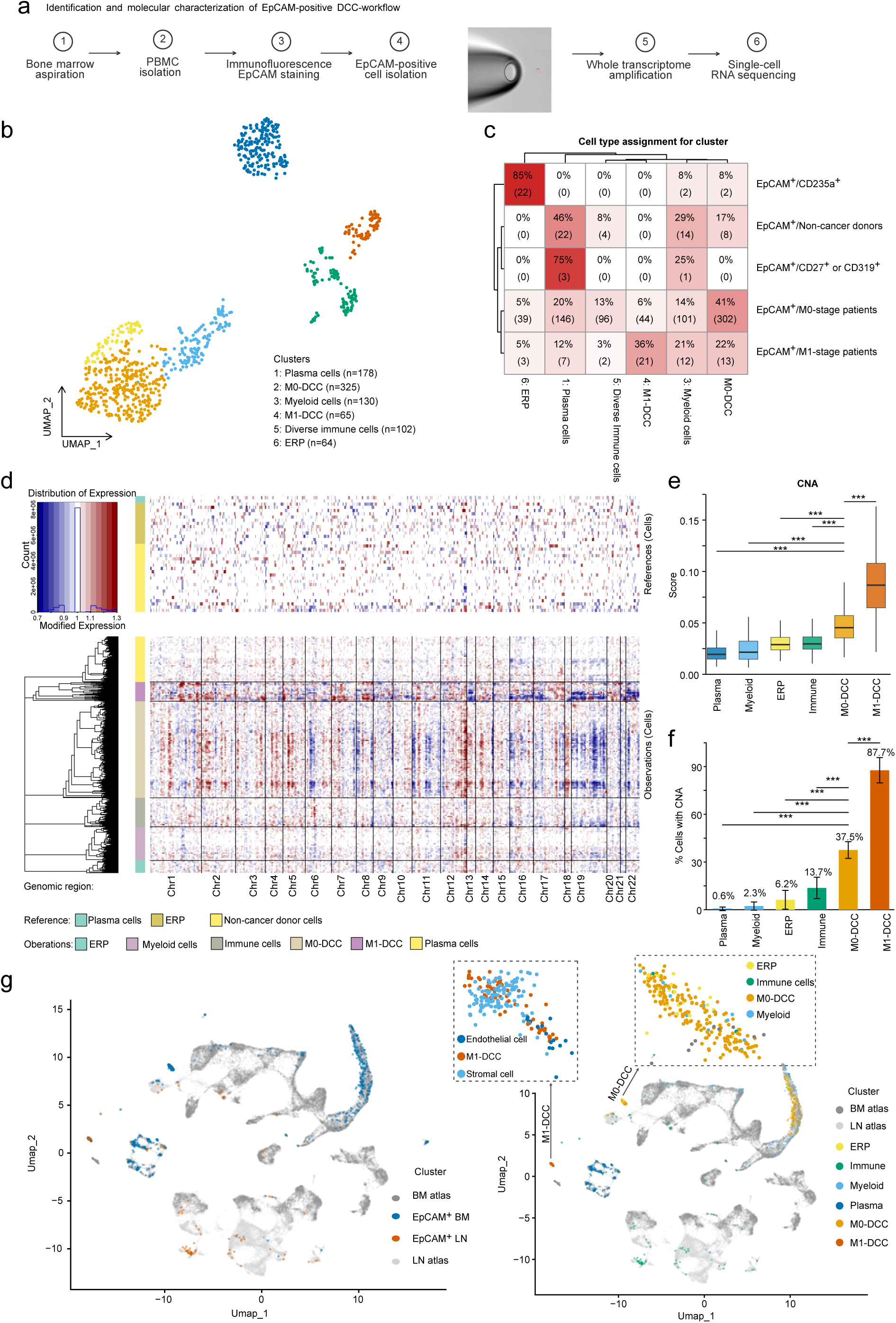
Detection, isolation and characterization of EpCAM+ cells from bone marrow and lymph nodes. **a) Workflow and cell isolation.** Left: Overview on sample preparation. For LN samples an additional step of disaggregation is needed. Right: Isolation of single EpCAM^+^ cell by micromanipulation **b) UMAP cluster**. Dimensionality reduction plot showing the clustering of EpCAM+ cells based on their transcriptional profiles. Each dot represents an individual cell, and colours denote the assigned cluster identities (plasma cells, myeloid cells, diverse immune cells, ERP, M0-DCC, M1-DCC), as defined in Fig 1c. **c) Cell composition of UMAP-Seurat clusters as defined by input**. The heatmap displays the percentage distribution of cells from each annotated cluster (columns) across different sample groups or conditions (rows). Values indicate the proportion of cells within each cluster assigned to each group, with the total cell count shown in parentheses. Targeted isolation of EpCAM^+^/CD235a^+^ and EpCAM^+^/CD319^+^ or CD27^+^ cells defined cluster 6 and cluster 1 as ERP and plasma cell cluster, respectively. Patient disease stage defined cluster 2 (M0-stage) and cluster 4 (M1-stage) as most cells in these clusters had been isolated from patients in these stages. EpCAM+ cells from non-cancer donors were distributed into various clusters. The color gradient represents the percentage, with darker shades indicating higher proportions. **d) InferCNV heatmap of copy number alterations across cell clusters.** Columns represent genomic regions (chromosomes 1–22), and rows represent individual cell clusters (excluding reference cells in lower panel) or reference cells. M0-DCC and M1-DCC clusters show a predominantly abnormal copy number state across the genome. **e) Distribution of CNA scores across cell clusters.** Box plot of CNA scores for each annotated cluster, with higher scores corresponding to greater genomic instability or larger deviations from the diploid state. Wilcoxon rank-sum tests (Mann-Whitney U tests) were performed to compare CNA scores (MAD values) between the M0 cluster and each of the other cell clusters. P-values were adjusted using the Benjamini-Hochberg (BH) method correction to control the expected proportion of false positives. *** indicate p-adj < 0.001.Each box represents the interquartile range (IQR), with the horizontal line indicating the median. Whiskers extend to the most extreme values within 1.5 × IQR from the quartiles. **f) Proportion of cells with detectable CNA across clusters.** The bar plot displays the percentage of cells within each annotated cluster that harbour detectable CNAs. Clusters show varying levels of CNA prevalence, ranging from 0.6% in plasma cells to 87.7% in M1 cells, indicating differential genomic instability across populations. Error bars represent the 95% confidence interval for the proportion, calculated as p ± 1.96 × SE, where SE = sqrt(p × (100 - p) / n). Statistical significance of differences in CNA prevalence between M0 and each other cluster was assessed using two-sided Fisher’s exact tests, with p-values adjusted for multiple comparisons using the Benjamini-Hochberg (BH) method. *** indicate p-adj < 0.001. **g) Integration of EPCAM+ cells with BM and LN atlases.** Left panel: UMAP projection showing the integration of EPCAM^+^ cells from this study (coloured by tissue origin: BM or LN) with reference atlases from bone marrow (BM atlas) and lymph node (LN atlas). Most query cells correctly integrate with their tissue-of-origin counterparts from the atlases. Right panel: Cells are coloured by their assigned cluster identities or atlas. Zoom-in visualization shows that M1-DCC cluster with stromal and endothelial cells as most similar neighbours, whereas M0-DCC did not find matching and form a cluster of their own.

### EpCAM staining in BM identifies various cell types

We performed UMAP and graph-based cluster analyses ^19^ to assess the heterogeneity of EpCAM+ transcriptomes. This grouped the cells into six clusters (Fig 1B). To assign cluster identities, we used the input information to designate each cluster according to its predominant cell type or clinical stage (Fig 1C). For example, 85% of EpCAM+/CD235a+ cells belonged in cluster 6, which was consequently referred to as ‘ERP’ (Fig.1 B and C). CD27^+^ or CD319+/EpCAM+ cells defined plasma cell cluster 1. EpCAM+ cells from M0- and M1-stage patients contributed to all clusters; clusters 2 and 4 mostly comprised cells with no known identity and were respectively designated M0-DCC and M1-DCC. Cells from non-cancer controls were found in all clusters, except clusters 4 (M1-DCC) and 6 (ERP). We also obtained 137 EpCAM+ cells from LN of NSCLC patients, but they exhibited distinctive gene expression profiles compared to BM (Fig S1A-C). Cells from clusters 3 and 5 comprised myeloid and immune expression patterns (FigS1C), with immune cells mostly derived from LN. Many LN cells fell into M1-DCC cluster 4, albeit that almost all were derived from patients at clinical stage M0 (FigS1B). Accordingly, many LN-derived NSCLC EpCAM+ cells clustered with prostate and breast cancer DCC from patients with manifest metastasis, corresponding to M1-DCC.

### Inferred copy number alteration (CNA) suggests malignant origin of M0- and M1-DCC

We controlled for cluster stability by cluster consistency analysis (Fig S1D), performed differential gene expression between cell clusters (Fig S1E), and inferred copy number alteration (CNA) for individual cells in different clusters (Fig 1D). The highest proportion of cells with inferred CNAs were M0-DCC (38%; cluster 2) and M1-DCC (88%; cluster 4; Fig 1 D-F). ERP, plasma cells, and myeloid BM cells exhibited CNA in ≤6% of cells, immune cells <14%. Aberrant immune were mostly B cells known to be competent for genome rearrangements (Fig 1F). The inferred CNA increase from M0- to M1-stages was consistent with previous results assessing individual DCC DNA ^20–23^. When all BM and LN EpCAM+ cells were projected onto a single-cell atlas (Fig 1G), ERP, plasma cells and diverse myeloid and immune cells integrated into the expected cell areas, with mostly correct assignment of tissue origin: for example, BM plasma cells *vs* LN plasma cells (Fig 1G). M1-DCC cells co-clustered with BM stroma cells (Fig 1G, S1G), possibly because they reflected activation of the epithelial-mesenchymal transition (EMT; see below). In stark contrast, no neighbouring atlas-derived cell type was found for M0-DCC. M0-DCCs therefore display a gene expression profile with little or no similarity in human BM or LN (Fig 1G, S1G, H). Together, our analyses confirmed that, in patients with carcinomas, EpCAM staining identifies populations with two expression profiles (M0- and M1-DCC) that are distinct from EpCAM+ ERP, plasma cells, diverse myeloid and immune cells and from all other cells of these organs. M0- and M1-DCC gene expression correlates with the clinical stage of their respective cancers (*i.e*. with or without manifest metastasis) and they exhibit relative chromosomal instability.

### M0-DCC and M1-DCC display distinct gene expression profiles

Cells from clusters 2 and 4 (henceforth respectively M0-DCC and M1-DCC) were positioned far apart in the BM-LN atlas (Fig 1G), indicative of widely differing gene expression programs. This is fully consistent with previously identified genomic differences of M0- and M1-DCC ^3,7,23,24^. From NSCLC, prostate and breast cancer patients, we had isolated a sufficiently high number of M1-DCC to enable us to retrieve characteristic histogenetic marker genes from their gene expression profiles (Fig 2A). Applying this gene list to M0-DCC from the same cancer types (*eg* breast, prostate, lung) resulted in no clustering (Fig 2B), indicating that other genes would be necessary to discriminate between M0-DCC and M1-DCC. Only Male vs female origins were distinguishable from *XIST* expression (Fig 2B). M1- and M0-DCC differed in all categories investigated, including GO-categories, hallmarks, and cell type assignments (Fig 2C). For ERP, plasma, myeloid, and immune cells these assignment were fully consistent with their atlas integration (Fig S2A-D). A direct comparison of M0-DCC *vs* M1-DCC for enriched cancer-associated pathways revealed the expected terms for M1-DCC, including epithelial origin and EMT, whereas stemness-associated terms predominated M0-DCC (Fig 2D), consistent with the loss of many tissue-defining markers (Fig 2A, B). Because cancer cell heterogeneity has previously been mostly assigned to epithelial phenotype variation ^25^, we were surprised to find that M1- and M0-DCC (derived from epithelial cancers), differed for the expression of cancer metaprograms ^25^, with stress, EMT and epithelial senescence being the most characteristic for M1-DCC, and the non-epithelial red blood cell (RBCS), MYC-activation and cell cycle for M0-DCC (Fig S2E). This further supports the notion that M0-DCC and M1-DCC manifest fundamentally different genome expression profiles.

**Figure 2:**
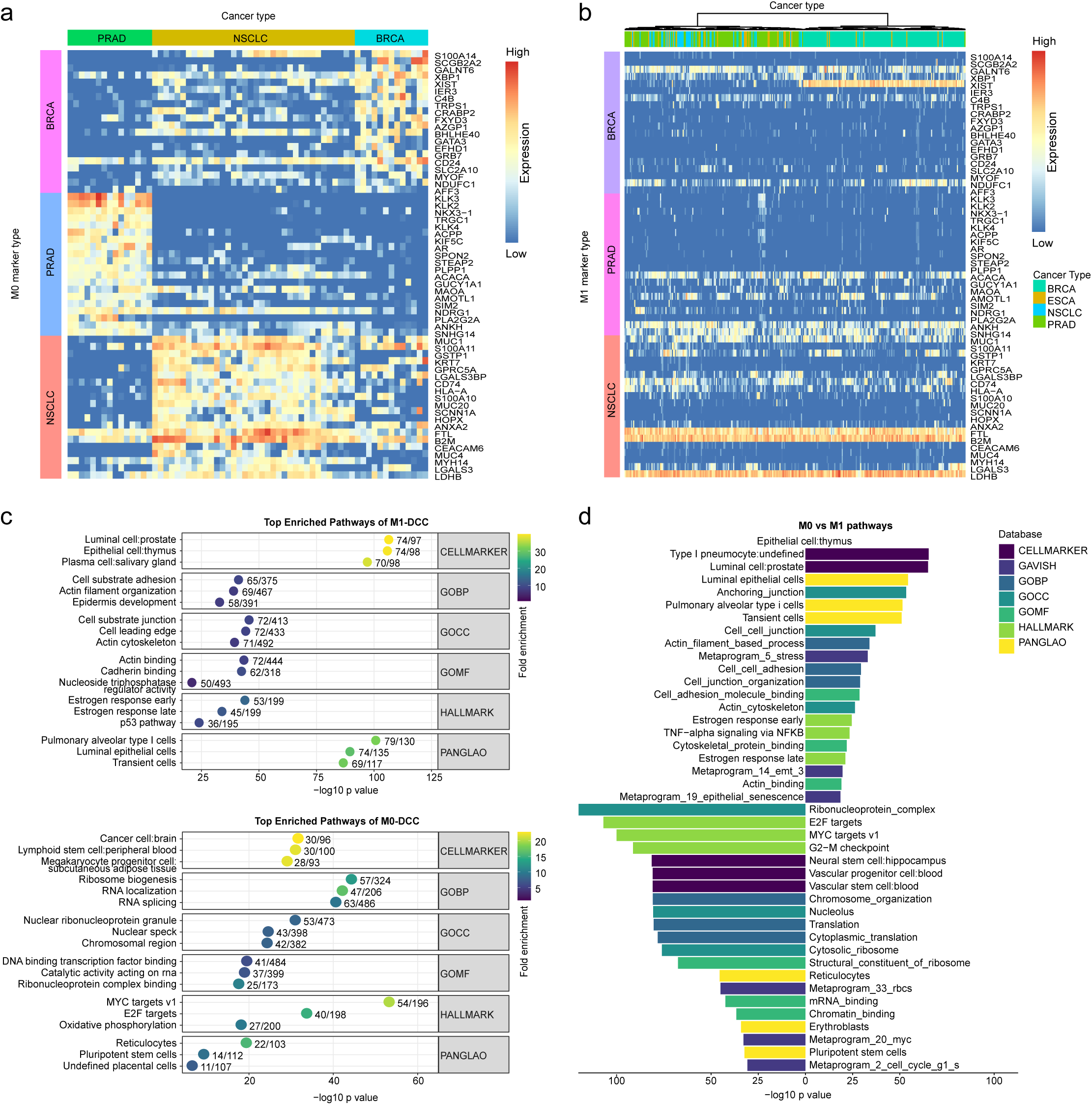
Comparison of M0-DCC and M1-DCC. **a) Histogenetic markers separate M1-DCC but not M0-DCC by tissue origin.** Heatmap displaying the marker gene expression of M1-DCC from different cancer types with sufficient cell numbers (BRCA, PRAD, NSCLC) using M1-DCC marker genes. Rows represent individual genes, and each column is one cell. Row labels display cancer type specific markers and column labels indicates cancer origin. Note the expression of typical markers for each cancer, such as KLK3 (PSA) for prostate, GATA3 for breast, and KRT7 for NSCLC. b) **No separation of M0-DCC by histogenetic marker genes.** Heatmap displaying the expression of the same marker genes as in panel a) across M0-DCC. Note the loss of KLK3, GATA3, CD24 and KRT7. Breast cancer-derived cells and female NSCLC or ESCA cells are only clustered by the expression of the non-coding RNA XIST, being expressed exclusively from the X-chromosome inactivation centre of the inactive X chromosome but not in male cells. c) **Dot plot showing the top enriched pathways identified in M1-DCC and M0-DCC.** Enrichment is shown the databases GOBP, GOMF, GOCC, HALLMARK, panglao and cellMarker. X axis displays the negative log10(p value) and colours indicate fold enrichment. Top three categories of each database are shown. d) **Direct comparison of pathway enrichment of M0- vs M1-DCC.** Bar plots showing the top enriched pathways identified for the two cell clusters. Bars are coloured by database origin. In addition to epithelial properties, M1-DCC are enriched in cell junction and cancer EMT, whereas M0-DCC are enriched for stemness features and the red blood cell metaprogram. Top three categories of each database are shown.

Although M0-DCC had largely inactivated gene expression programs associated with presumptive tissues-of-origin, we searched for residual gene expression corresponding to these origins. Cluster analysis indicated modest transcriptome differences between M0-DCC from different organs, indicating substantial erasure of tissue-of-origin gene expression (Fig 3A).

**Figure 3:**
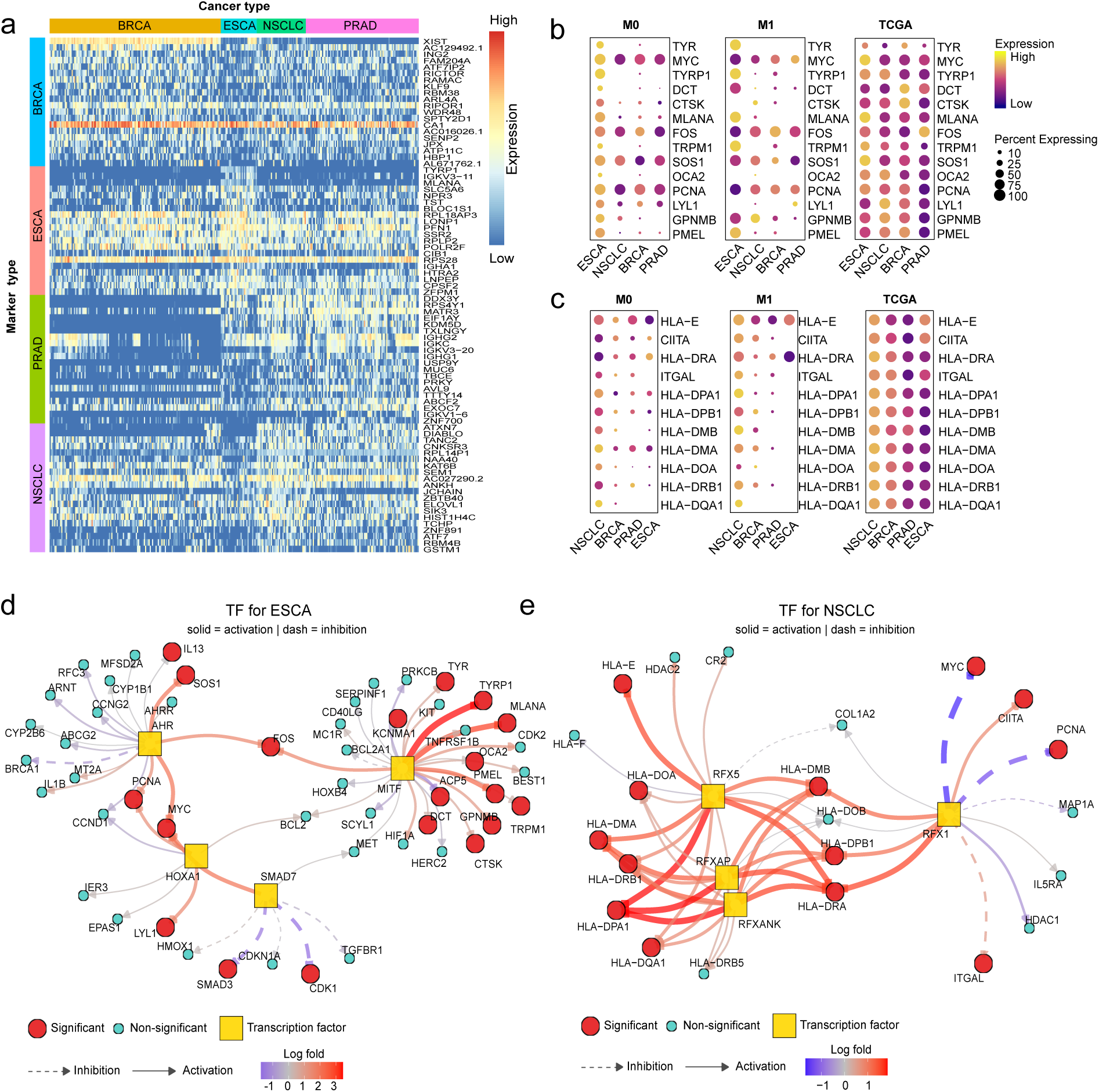
Traces of tissue origin in M0-DCC. **a) Heatmap displaying top 20 cancer type specific M0 marker genes across cells in the M0-DCC cluster.** Rows represent individual genes, and each column is one cell. Row labels indicate marker gene origin and column labels cancer origin. Compared to M1-DCC in Fig 2a, only a weak tendency of clustering is observed among M0-DCC. The most prominent signal is visible for ESCA. **b) and c) Detection of an underlying tissue specific gene expression network in ESCA (b) and NSCLC (c).** Dot plot showing the expression of M0-ESCA-DCC (oesophageal cancer) and M0-NSCLC-DCC-specific transcription factor target genes across M0-DCC, M1-DCC, and TCGA samples. Samples are grouped by tumour of origin. Dot size represents the percentage of samples expressing the target gene, and colour indicates the expression level. Target genes show a consistent expression pattern across M1-DCC, TCGA, and M0-DCC, confirming that M0-DCC cells originate from ESCA or NSCLC tumours, respectively. **d) and e) Network plot showing active transcription factors (TF) in ESCA and NSCLC**. Yellow squares indicate TFs that are significantly more activated in ESCA. Round nodes represent target genes of these TFs, where red node are genes that is significantly changed in ESCA M0 compared with other M0 cancer types and blue nodes are not significantly changed genes. Lines connect TFs to their target genes, with line type indicating whether the target gene is positively (solid line) or negatively (dashed line) regulated. Line color reflects differential expression in RNA-seq data (upregulated or downregulated).

To uncover previously undetectable cancer-type-characteristic gene expression, we performed transcription factor regulatory network analysis to confirm that predicted regulators and target genes were (i) characteristic for the subtype of M0-DCC, (ii) expressed by M1-DCC of the same subtype and (iii) expressed in The Cancer Genome Atlas (TCGA) data for the corresponding tumour type. Genes compliant with all three categories should be cancer-type associated. This was readily achieved for oesophageal cancer (ESCA) and NSCLC (Fig 3B and C), which possessed structured underlying TFRNs (Fig 3D and E). Sex-associated regulators and transcription factors dominated the terms in prostate and breast cancer cells, but we were unable to deduce clear signals of tissue origin.

In sum, whilst M1-DCC resemble cancer cells of well-known and expected phenotypes, M0-DCC display mere traces of the gene expression programs characteristic for their putatively corresponding tissue-of-origin. We next explored their underlying gene expression programs at higher resolution.

### M0-DCC gene expression resembles that of cleavage-stage, preimplantation embryos

Direct comparison between M0- and M1-DCC indicated greater stemness in M0-DCC (Fig 2D). We therefore sought deeper stemness and potency characterization using cytotrace, fitdevo and SC_mRNAsi, which most accurately assigned expression profiles to fetal or adult origin (Fig S3A). When applied to all EpCAM+ cells from BM and LN, the highest scores were obtained for M0-DCC (Fig 4A), exceeding even ERP, which otherwise ranked highest in adults. When all EpCAM+ cells were projected onto an atlas comprising the human cell landscape (HCL) and the BM atlas, M0-DCC mapped to fetus-derived cells or fetal or adult ERP, *i.e*. areas with highest stemness and potency scores (Fig 4B, Fig S3B). M1-DCC and all other cell types projected onto areas from adult tissues (Fig 4B). At higher resolution, CNA-containing M0-DCC (Fig S3B) mapped to a region corresponding to human embryonic stem cells (Fig 4B), and predominantly to primordial germ cells and fetal epithelial progenitor cells (Fig 4C).

**Figure 4:**
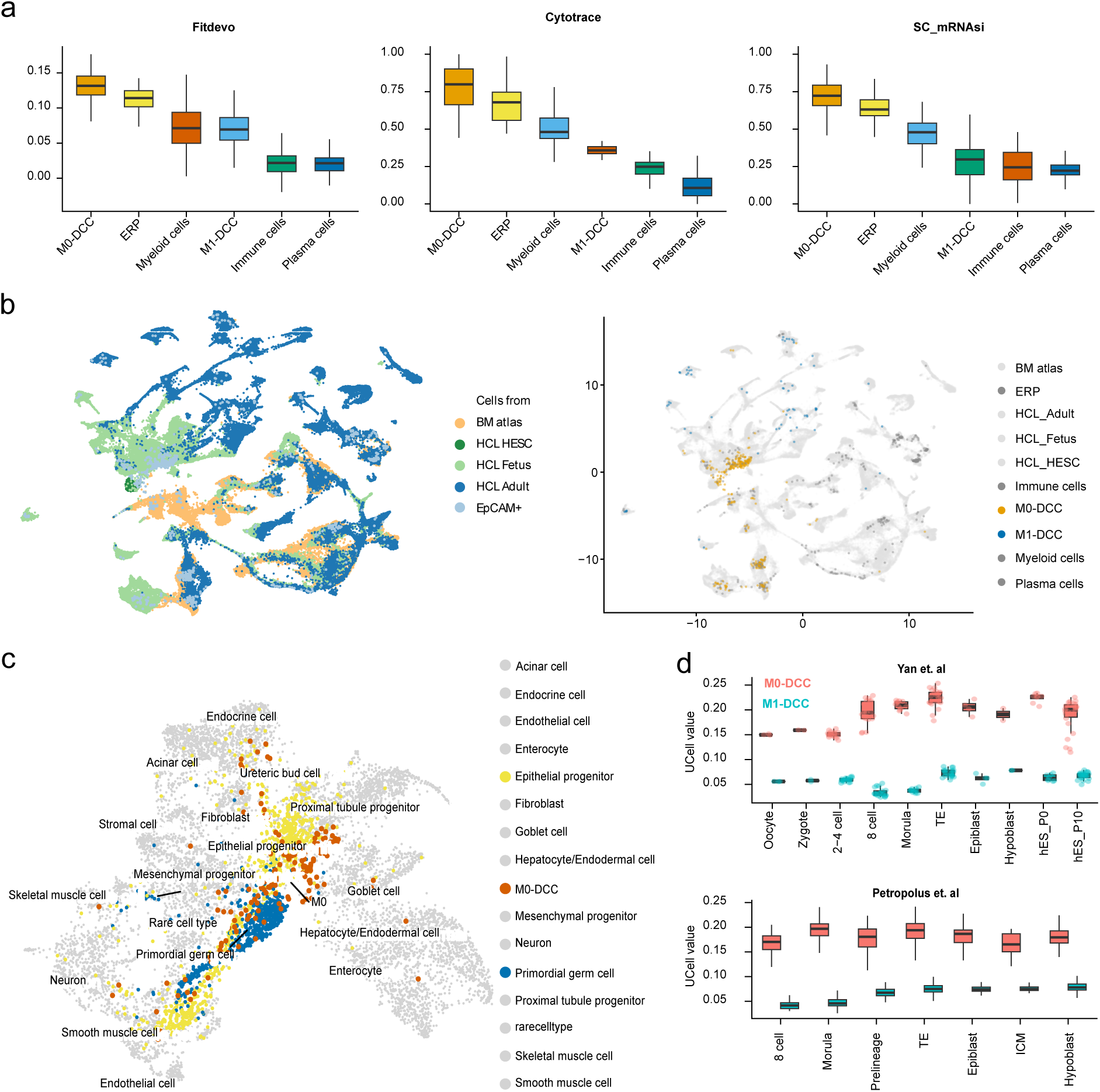
M0-DCC show striking parallels to early embryos. **a) Stemness and potency scores across EpCAM^+^ cell clusters assessed by three independent methods**. Box plots showing the distribution of stemness scores across six cell clusters in descending order (M0-DCC, ERP, M1-DCC, myeloid, diverse immune and plasma cells), calculated using three different stemness assessment tools (fitdevo ^51^, CytoTRACE ^50^, and SC_mRNAsi ^52^). Each box represents the interquartile range (IQR), with the horizontal line indicating the median. Whiskers extend to the most extreme values within 1.5 × IQR from the quartiles. **b) Integration of EpCAM+ cells into an extended human cell atlas comprising fetal tissues.** Left panel: UMAP projection showing the integration of EPCAM^+^ cells of Fig 1B into a combined BM and Human Cell Landscape (HCL) atlas, which includes HCL_HESC (human embryonic stem cells), HCL_Fetus (fetal tissues), and HCL_Adult (adult tissues). Each dot represents a single cell coloured by origin. Right UMAP panel: Cells are coloured by cluster identity (Plasma, Myeloid, ERP, Immune, M0-DCC, M1-DCC). M0-DCC cells predominantly integrate with HCL_Fetus, while M1-DCC cells show diverse integration across HCL_Adult. Note that other EPCAM^+^ cells integrate into regions of the BM and HCL comprising adult cells, demonstrating that fetal / embryonic integration is unique to M0-DCC. **c) UMAP projection of the region containing the majority of M0-DCC cells**. Cells are coloured by cell type annotation. M0-DCC cells predominantly co-localize with fetal epithelial progenitor and primordial germ cells. **d) Characteristic M0-DCC genes are highly expressed in early embryos.** Box plots showing the distribution of UCell scores for M0-DCC and M1-DCC marker gene sets in human embryonic cell types spanning developmental stages from oocyte to hypoblast ^27,28^. Higher UCell scores indicate stronger enrichment of the respective DCC cluster signature. The box represents the IQR, the centre line denotes the median, and whiskers extend to 1.5 × IQR.

M1-DCC transcriptome profiles shared the features of EMT, which reflects implantation and gastrulation in embryo development ^26^, leading us to assess whether M0-DCC represented cells from the same or different embryonic stages. M0-DCC gene expression was compared with previously-determined early embryonic transcriptomes ^27,28^ from the one-cell embryo stage (during and after fertilization), through cleavage stages (one- to 16-cell embryos), up to hypoblast (primitive endoderm) formation in blastocysts ^29^. We first tested the expression marker genes in early embryos. M0-DCC and ERP genes were highly expressed throughout preimplantation embryos unlike M1-DCC and plasma cells markers (Fig S3 C, D). Consequently, U-cell scores of early embryos differed strongly for M0-DCC and M1-DCC signature genes throughout early embryonic development (Fig 4D) and upregulated M0-DCC genes were highly enriched in preimplantation embryos as opposed to M1-DCC genes (Fig S3E). Values for M0-DCC exceeded those for ERP from one-cell to morula stages (following compaction at the 16-cell stage, ∼3.5 days post-fertilization in humans) of preimplantation development and equilibrated thereafter (Fig S3F). These stages occur before the earliest manifestation of embryonic EMT during human implantation (6∼10 d post-fertilization) or gastrulation (14∼21 d post-fertilization) ^29^. In sum, M0-DCC display a higher similarity to preimplantation, cleavage-stage embryos than is known for any cell type in healthy adults and from now on, we refer to them as embryo-like (EL) DCC.

### Induction of embryo-like DCC gene expression

We have shown that in human clinical contexts and mouse models, metastasis is an early event, and that some DCC capture the earliest observable stage not only of metastasis but of cancer ^8,20,30,31^. Prior to clinically manifest metastasis, DCC are difficult to detect and occur in very low numbers. Therefore, direct characterization is technically limited, and expansion *in vitro* or *in vivo* has failed or produced only anecdotal reports ^9,32,33^. Hypothesizing that the M0-DCC population is transient (because it seems to be replaced in M1-stage patients) and results from cell plasticity, we attempted to induce an M0-like state in cancer cell lines to generate M0-DCC models. We first classified all CCLE cell lines by their M0-DCC and M1-DCC gene expression scores and selected M0- and M1-DCC-like for further studies (Fig S4A). We also hypothesized that the phenotype of M1-DCC-like cell lines could be shifted towards a M0-like phenotype.

Coculture of MCF-7 (human breast adenocarcinoma) cells with BM samples from three non-cancer donors did not induce a M0-DCC gene expression signature (Fig 5A, B). We selected six marker genes (*SLC22A16*, *SLC43A1*, *CPVL*, *KCNH2*, *CD36*, and *KIT*), characteristically expressed in EL-DCC, ERP and early embryos to monitor (by qPCR) upregulation of gene modules (Fig S4B). Interestingly, *EpCAM*, *CD36*, *KIT* were fully co-regulated until the morula stage (Fig S4B). Given the activation of the RBCS (red blood cells) metaprogram by EL-DCC (Fig 2D, Fig S2E), we further evaluated whether upregulation of the erythroid and embryonic gene modules might be linked to the erythroblastic island (EBI) that is formed by erythroid island macrophages (EIM) to engender the specific niche for ERPs ^34,35^. There are few reports of human EIM isolation ^36^, but three cytokines (IL33, ANGPL7, SERPINB2) have been shown to program erythroblasts ^37^. In addition to EBI cytokines, we tested IL6-trans-signalling, as IL-6 was previously identified as a regulator of a stem-like state in mammary epithelial cells and DCC and abundantly employed in BM 8. We first treated six cell lines (MCF-7, BT474, T47D, SkBr3, MDA-MB-231, and MDA-MB-231 BM-adapted subline MDA-MB-231-1833) with different combinations of the three EIM cytokines in addition to hyper-IL-6 (HIL6) to activate IL6-trans-signaling selectively38. Following treatment for four days, we observed upregulation of one or several genes of the erythroid-embryonic signature in most of the cell lines tested (Fig S4C). M1-DCC-like cell lines MCF7, SkBR3, T47D, and BT474 underwent pronounced upregulation of erythroid genes, while MDA-MB-231 and MDA-MB-231-1833 cell lines, which exhibit low M1-DCC scores and positive M0-DCC scores, responded weakly (Fig S4C). Notably, the metastasis-derived MDA-MB-231-1833 exhibited higher baseline expression of erythroid marker genes than its parental progenitor (Fig S4D). Individual transcripts across the cell lines were differentially responsive to the cytokines, with highest induction by all four cytokines combined.

**Figure 5:**
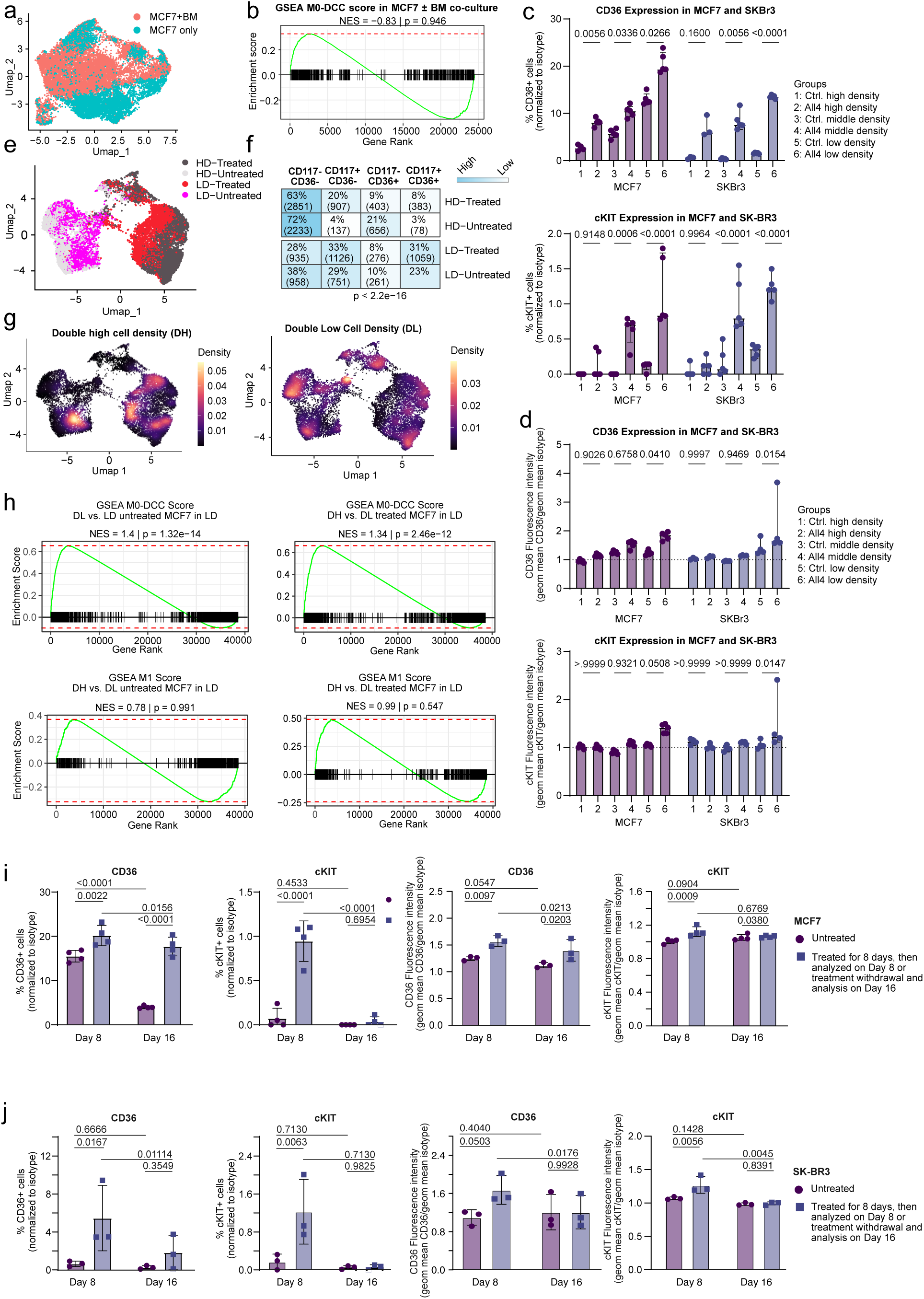
The embryo-like gene expression profile can be induced in cancer cell lines. **a) Coculture of MCF7 cells with human bone marrow does not induce the M0-DCC signature.** UMAP projection of MCF7 cells cultured alone or together with human bone marrow cells. **b) Gene Set Enrichment Analysis (GSEA) plot.** M0-DCC marker genes are plotted in a gene list ranked by a combined metric of fold change and statistical significance (log10(p-value) × sign of fold change) for the two conditions. Genes are ranked from high to low based on this combined score. The enrichment plot indicates that BM culture does not significantly shift MCF7 cells toward the M0-DCC transcriptional phenotype. **c) and d) Impact of cell density and BM cytokines on CD36 and KIT protein expression**. CD36 (top) and KIT (bottom) expression as percentage of positive cells (c) or expression intensity under different conditions for the breast cancer cell lines SKBr3 and MCF7. **e-g) Induction of the M0-DCC mRNA signature in single cells. e)** UMAP plot (top left) showing single cells coloured by sample condition, i.e. high (HD) vs low (LD) cell density, with and without cytokines (treated vs untreated). Each dot represents an individual cell. **f)** Heat map (top right) displays the percentage and absolute numbers of cells expressing high or low CD36 or KIT/CD117 as defined by the antibody DNA tag (ADT) across experimental conditions. A chi-square test p-value of 2.2×10⁻¹⁶ indicates a highly significant association between ADT status and samples, revealing that particularly LD-treated samples are enriched in double high cells. **g)** UMAP projections displaying the localization of DH (double-high) and DL (double low) CD36 or KIT/CD117 cells across samples defined by ADT. Each plot represents the local cell density, with brighter colours indicating higher density. **h) Induction of the M0-DCC gene signature in CD36^high^/KIT^high^ MCF7 cells.** Gene Set Enrichment Analysis (GSEA) plot testing the enrichment of M0-DCC and M1-DCC marker genes in gene lists ranked by double high CD36/CD117 (DH) vs double low CD36/CD117 (DL) in LD with and without cytokine treatment. Genes are ranked from high to low (DH vs DL). Top: M0-DCC markers show enrichment in DH vs DL cells with and without treatment, making DH cells more M0-DCC-like compared to DL cells. Bottom: no change in M1-DCC marker gene expression under the applied conditions. **i and j) Reversible CD117/CD36 expression after withdrawal of inducing cytokines.** The surrogate markers for the EL-DCC phenotype (CD36, CD117) return to baseline levels after withdrawal of inducing cytokines.

We tested whether mRNA expression correlated with increases in corresponding protein levels for CD36 and KIT/CD117. Since early DCC differ from manifest metastases in their lack of homotypic cell-cell contacts, we explored the combined effect of all four cytokines on MCF7 and SKBr3 (human breast adenocarcinoma) cells plated at different densities. Highest CD36 and KIT upregulation occurred at the lowest plating density of cells exposed to all four cytokines, albeit at low percentages (1-20%; Fig 5C, D). Plating density impacted CD36 or KIT upregulation in MCF7, but not SKBr3 (Fig 5C, D). Analogous treatments on two cell lines with low or negative M1-DCC scores (CAL51, DU4475) did not induce CD36 and KIT protein expression (Fig S4E).

These data were corroborated and extended by coupled scRNA-seq and CD36 and KIT protein measurement via antibody-DNA tag (ADT) oligonucleotide barcoding. Treatment of MCF7 cells with all four BM cytokines induced marked gene expression changes, especially at low plating densities (Fig 5E, Fig S4F, and strongly induced CD36^+^CD117^+^ expression (Fig 5F, G). We found a significant (p = 2.46 x 10^−12^ to 1.32 ×10^−14^) enrichment of the M0-DCC gene expression profile, but not that of M1-DCC, in strongly CD36^+^CD117^+^ cells compared to CD36^low^CD117^low^ cells (Fig 5H). Finally, we determined whether the cytokine-induced state was reversible. Removal of cytokines after eight days of induction resulted in an almost complete reinstatement of CD36 and CD117 starting values within eight days (Fig 5I, J).

In sum, an EL-DCC gene expression pattern can be reversibly induced in epithelial cancer cells by BM cytokines, dependent on microenvironmental signals.

### Clinical and functional consequences of embryo-DCC similarities

Some of the cell lines we evaluated exhibited high EL-DCC gene expression scores, whereas others more closely resembled M1-DCC gene expression. We therefore interrogated the CCLE data base ^39^ for a functional link between the patient-derived M0-DCC signatures and the formation of metastases in mice. Here, intracardiac cell injections into immunodeficient mice had been performed and metastasis was quantified in five organs (brain, bone, liver, lung, kidney) for 481 human cell lines, representing 21 solid tumour types ^40^. Provenance grouping into primary tumours (n=286) and metastases (n= 176) revealed that cell lines from metastases generated significantly (p=0.00028) more metastases (Fig 6A). M0-DCC^high^ cell lines (n=190) were associated with increased metastasis rates with even higher significance (p= 2.9 ×10^−6^), contrasting with cells whose expression patterns more closely resembled those of M1-DCC (n=293). Here, negative M1-DCC scores were linked to metastasis formation in mice, whereas positive M1-scores seemed to protect against metastasis initiation (Fig 6A). Since M1-DCC are from patients with manifest metastasis, these findings suggest that metastatic initiation utilizes a genetic program distinct from metastatic expansion.

**Figure 6:**
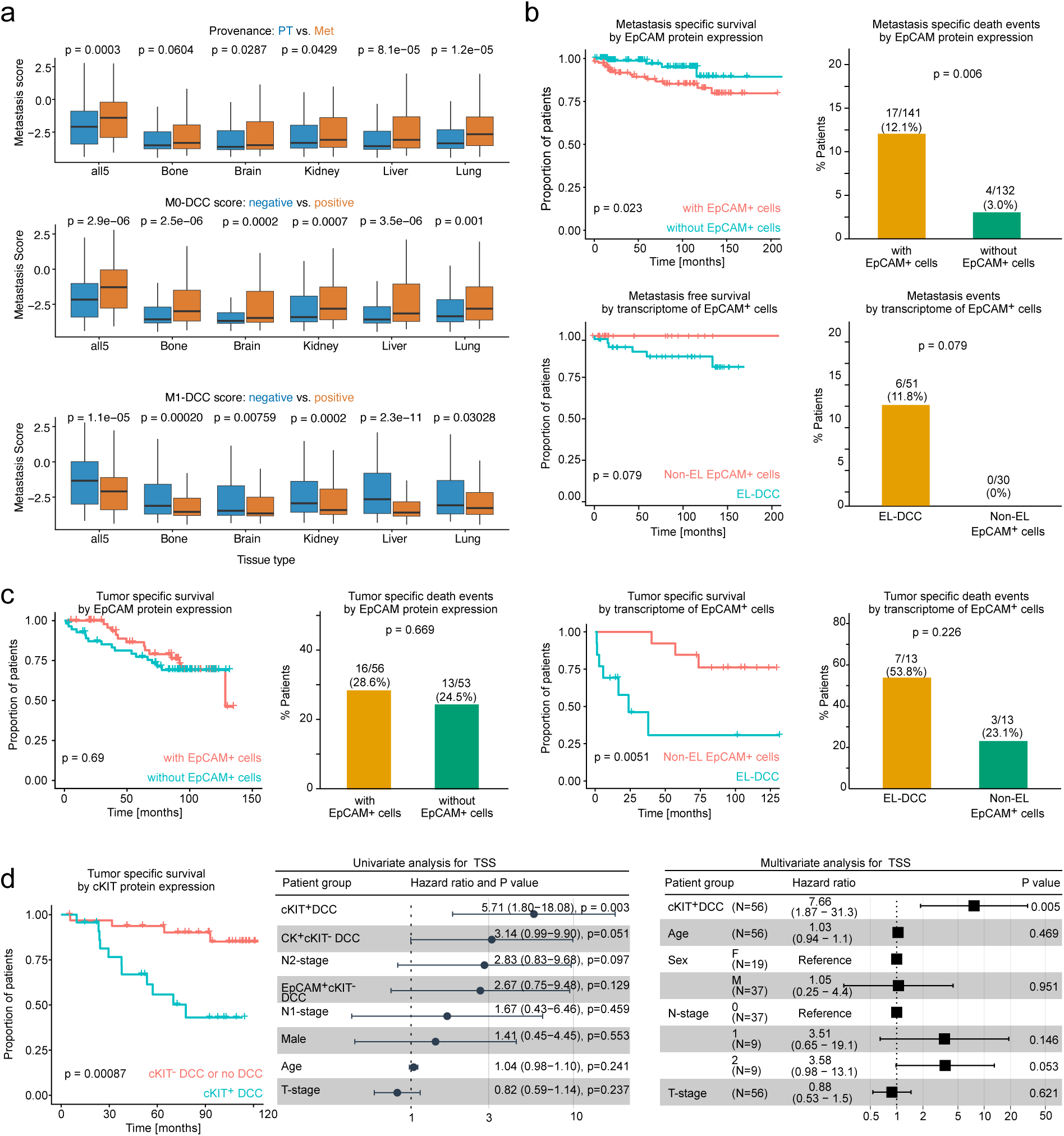
Impact of the M0-DCC phenotype on *in vivo* metastasis and patient outcome. **a) High M0-DCC scores predict metastasis of human cell lines to all major sites.** Box plot showing metastasis scores of cancer cell lines grouped either by origin (primary tumour derived vs metastasis derived; top panel); by their M0-DCC marker gene set z-score (positive vs negative); by their M1-DCC marker gene set z-score (positive vs negative). Statistical significance was assessed using Wilcoxon rank-sum test. Note that the M1-DCC signature, representing cells from manifest metastases, is associated with reduced metastatic potential, suggesting different roles for M0-DCC and M1-DCC transcriptional programs in cancer metastasis. **b) and c) Impact of embryo-like EpCAM+ cells on breast and lung cancer outcome.** Follow-up data are currently available for breast and lung cancer patients. b) Kaplan-Meier curves for metastasis-free survival (MFS) of BRCA (breast cancer) M0 patients grouped by EPCAM staining status (positive vs. negative) and by presence of embryo-like EpCAM+ cells among EpCAM+ cells in BM. Bar plots display the proportion of BRCA (breast cancer) patients with and without metastasis for both comparisons. c) same analysis for tumour-specific survival in NSCLC patients. **d) Protein expression of KIT by DCC identifies patients at high risk of cancer death.** Kaplan-Meier curve showing tumour specific survival (TSS) of NSCLC M0-stage patients grouped by KIT expression on cytokeratin^+^ or EpCAM^+^ DCC in BM (left panel). Forest plots showing univariate (middle panel) and multivariate (right panel) Cox regression analyses for survival prediction in NSCLC M0-stage patients. Univariate analysis included KIT staining, CK staining, EPCAM staining, TNM T grade, N status, Sex, and Gender. Only patients with all three staining markers (KIT, CK, EPCAM) were included. Multivariate analysis included KIT staining (the only significant factor in univariate analysis) along with conventional clinical annotations (T stage, N stage, sex, and age).

Harnessing prospective data for breast cancer and NSCLC patients from whom we had isolated BM EpCAM^+^ cells, we noted a clear impact of EL-DCC (Figure 6B-D). Breast cancer patients with EpCAM^+^ cells had reduced metastasis-free survival compared to patients without detected EpCAM^+^ cells (p= 0.023). Of note, in patients with EpCAM^+^ cells, all metastatic events occurred in the group in which EL-DCC had been detected (Fig 6B). For NSCLC patients, tumour-specific survival was similar for patients with and without EpCAM^+^ DCCs, suggesting that EpCAM staining missed patients dying from the disease (Fig 6C). However, when EpCAM^+^ patients were stratified according to cluster assignment, those with EL-DCC had very poor tumour-specific survival (Fig 6C). To overcome the limitation of EpCAM as single DCC marker in NSCLC and to validate the EL-DCC phenotype, we screened archived BM samples from 56 NSCLC patients by cytokeratin (CK), EpCAM and KIT triple staining. We found that curatively-treated M0-stage patients with CK^+^KIT^+^ or EpCAM^+^KIT^+^ or CK^+^EpCAM^+^KIT^+^ DCC (i.e. KIT^+^DCC) had the highest risk of progression (Fig 6D): multivariable analysis indicated that a single KIT^+^DCC in BM outcompeted all clinical risk factors (Fig 6D).

## Discussion

We here describe an inferred cellular phenotype, previously unknown in adults. Positive EL-DCC gene expression signature scores in cell lines is associated with a high propensity to initiate metastasis in all tested organs upon intracardiac injection and predicts metachronous metastasis. Detection of EL-DCC in bone marrow predicts very poor outcome in breast and NSCLC patients. Strikingly, their gene expression resembled that of cleavage-stage preimplantation embryos and is therefore clearly distinct from EMT, a process whose cancer association with later-stage implanting and gastrulating embryos is well-established ^26,41^. Apparently, parallels between cancer and early development are closer than previously thought. We therefore suggest the term embryo-like DCC (EL-DCC) to describe DCCs that recapitulate cleavage-stage embryonic gene expression, and the term metastasis founder cell (MFC) for their associated function. EL-DCC gene expression is inducible by BM-derived cytokines, specifically cytokines from EIM, a very rare macrophage subtype controlling haematopoiesis ^37^. The overlap between EL-DCC and ERP gene expression programs seems related to their high stemness scores and possibly to the exposure and responsiveness to EIM cytokines.

The identification of EL-DCC is likely to have significant consequences for metastasis research. First, early systemic cancer (M0-stage) and manifest metastasis (M1-stage) are not only anatomically distinct disease stages, but the target cells of systemic therapies in these stage are fundamentally different - so different that prevention of metastasis will likely require novel approaches. Furthermore, EL-DCC may turn out to provide excellent therapeutic markers and targets, informing pharmacological strategies to prevent metastasis, thereby significantly shortening (neo)adjuvant therapy studies. The potentially unique gene expression profile of EL-DCC may reveal therapy targets and biomarkers that are absent in adult patients, with consequences for early detection, monitoring and drug development. We are aware that the presence of EL-DCC raises more questions than it answers but their discovery promises to open novel approaches to removing the Damocles’ sword of metastatic relapse that today threatens millions of patients.

Finally, the close relatedness of EL-DCC and cleavage-stage embryogenesis holds the remarkable potential to illuminate mechanisms of cancer initiation. With the exception of heritable cancers - a small proportion (5-10%) of the total ^42^ - little is known about how cancer initiates, largely because the initiating events have been over-written by the time disease is overt. By contrast, the process that triggers embryogenesis is known: fertilization. The echoing of transcriptional regulation in preimplantation development by transcription in EL-DCCs also chimes, with the onset of embryonic transcription (immediate embryonic genome activation, iEGA) during and after fertilization: iEGA implicates oncogenic regulators, including MYC ^43–45^ some of which are implicated in EL-DCC. This work accordingly provides strong evidence that the onset of cancer recapitulates fertilization, at least in part. To the extent that this is correct, it has disruptive implications for efforts to uncover mechanisms underlying the onset of some cancer and the detection and treatment of disease before it is clinically manifest.

## Acknowledgements

We thank Anthea Povall, Isabell Blochberger, Justin Skotnitzki, Nathalie Drexler, Sophie Hochmuth, Irene Nebeja, Stefanie Güldener for excellent technical assistance. We thank all technicians, scientists and students who helped to screen bone marrow samples for EpCAM^+^ cells, namely Thomas Schamberger, Manfred Meyer, Nina Patwary, Gundula Haunschild, Lisa Maria Köhler, Susanna Lissek, Laura Rudhart, Hans-Jürgen Laberer. This work was supported by grants to CAK from the Deutsche Forschungsgemeinschaft (DFG; KL 1233/10-1 and 2, /11-1 and 2, /12-1, /18-1, /19-1, TRR 305 Z02 and A01), Deutsche Krebshilfe (111536), the ERC (ERC-2012-ADG-322602); a grant from HiberCell to Fraunhofer ITEM-Regensburg, from the DFG to MG GU 1923/1-1 and 2).

## Authors’ Contributions

**Conception and design:** C. A. Klein

**Development of methodology:** H. Koerkel-Qu, Elia Raya, C. A. Klein

**Acquisition of data or samples:** Elia Raya, M. Guzvic, C. Irlbeck, T. Mederer, D. Spitzl, Z. Czyz, L. Schunicht, S. Seitz, J. Roth, B. Rack, H. Kurdieh, N. Harbeck, R. Mayr, M. Burger, T. Robold, H.-S. Hofmann, M. Weber, K-P Janssen, S. Hücker, S. Kirsch, M. Werner-Klein

**Analysis and interpretation of data:** H. Koerkel-Qu, Elia Raya, M. Guzvic, T. Mederer, D. Spitzl, L. Schunicht, S. Kirsch, S. Hücker, A.F.C. Perry, C. A. Klein

**Writing of the manuscript:** H. Koerkel-Qu, E. Raya, A.F.C. Perry and C. A. Klein

**Review and/or revision of the manuscript:** all authors

## Competing interest

H. Koerkel-Qu, E. Raya, M. Werner-Klein, T. Mederer, D. Spitzl, M. Guzvic, C. A. Klein are inventors on an IP application of the Fraunhofer Society.

## Methods

### Patient inclusion and ethics statement

The study complied with all relevant ethical regulations governing the use of human material. For patients with non-small-cell lung cancer (NSCLC), bone marrow (BM) aspirates and lymph node (LN) samples from the first and second lymph node stations were collected during potentially curative surgery for suspected or histologically confirmed NSCLC at the Department of Thoracic Surgery, University Hospital Regensburg, or at Hospital Barmherzige Brüder Regensburg. BM aspirates were additionally obtained from patients with breast cancer (BRCA) treated at LMU Munich or Caritas-Krankenhaus St. Josef Regensburg, from patients with prostate cancer (PRAD) treated at Caritas-Krankenhaus St. Josef Regensburg and from patients treated oesophageal cancer Technical University of Munich. The breast and prostate cancer cohorts included patients both without (M0-stage) and with clinically detectable distant metastases (M1-stage). EpCAM+ cells were also obtained from BM of patients without known malignant disease undergoing hip replacement or trauma surgery (non-cancer donors; University hospital Regensburg, Krankenhaus Barmherzige Brüder Regensburg, Klinikum Dachau, Klinikum Bad Abbach). Written informed consent of cancer and control patients was obtained and the ethics committee of the University of Regensburg (ethics vote numbers 07/79 and 18-948-101) enabling tissue sampling and analysis of isolated cells.

### LN and BM processing

The lymphatic tissue was cut into 1-mm pieces and disaggregated mechanically into a single-cell suspension by rotating knives (DAKO Medimachine, DAKO), washed with HBSS (Life Technologies, Heidelberg, Germany) and centrifuged on a density gradient made of a 60% Percoll solution (Amersham, Uppsala, Sweden). Bone marrow samples were twice washed with Hank’s balanced salt solution to remove fat and thrombocytes. Next, cell suspension was centrifuged in density gradient made of a 65% Percoll solution (Amersham, Uppsala, Sweden). After centrifugation, the interphase containing mononuclear cells (MNC) were carefully collected and washed with PBS. The number of MNCs and erythrocytes was determined on hemocytometer. To enrich the DCC-containing fraction, the sample was depleted of most hematopoietic cells using negative immunomagnetic selection. This was achieved by incubating the cell suspension with APC-conjugated antibodies against CD11b, CD33, and CD45. After incubation and washing, the cell suspension was incubated with anti-APC beads and anti-CD235a beads (glycophorin A; all Miltenyi). After incubation and washing, the cell suspension was run through the 40-μm cell sieve and then run on LS MACS column. The eluate, containing the unlabeled cell fraction, was collected on ice and the cell number determined using a hemocytometer.

### Staining and screening of BM and LN samples

On average, two million BM or LN cells were stained with anti-EpCAM-PE (HEA125, Miltenyi Biotec) antibody. Each sample was manually screened for the presence of EpCAM^+^ cells on an inverted fluorescent microscope (Olympus or Zeiss), equipped with micromanipulator (Patchman NP2, Eppendorf) and pump (CellTram, Eppendorf). Single cells with preserved morphological integrity and positively stained for EpCAM were extracted using a glass capillary attached to the micromanipulator. By visual inspection, we ensured that only one cell was in the capillary. The cell was transferred to an empty field with PBS and manually isolated using a micropipette. After isolation of single EpCAM^+^ cells from each sample we isolated a pool of approximately 2,000 to 3,000 cells. As a reagent control for contaminating nucleic acids, 1 μL of the PBS, in which individual EpCAM^+^ cells were isolated, was taken for subsequent whole transcriptome amplification (WTA).

### Whole transcriptome amplification (WTA)

Messenger RNA isolation from single cells, reverse transcription and global amplification of first-strand complementary DNA were performed as described previously ^4,46^. WTA product quality was assessed by a multiplex endpoint PCR assessing the presence of three housekeeping gene ^9^. High quality was assigned to cells with at least one of three transcripts detected.

### Next-generation sequencing of single-cell mRNA (scRNA-seq) library preparation and sequencing

Single cell RNA-seq libraries were prepared were prepared from individual EpCAM+ cells isolated from patients with cancer and non-cancer controls, as previously described ^9^. Libraries were first subjected to shallow sequencing on an Illumina MiSeq System (MiSeq Reagent Kit v2, 50 cycles) for library quantification, pooled at equimolar ratios, and subjected to deep sequencing on an Illumina NovaSeq 6000 platform.

### scRNA-seq data analysis

Reads were aligned to the human reference genome (GRCh38) using STAR (v2.6.1c), and gene-level expected counts were quantified using RSEM (v1.3.1). Low-quality cells were removed based on the following quality control (QC) criteria: library size <50,000 reads, detected genes< 1000, or > 50% mitochondrial gene counts. Doublets were computationally identified and excluded using scDblFinder (v1.26.1). Cells passing filtering were normalized by calculating size factors using computeSumFactors from scran (v1.40.0), followed by log-transformation via logNormCounts from scuttle (v1.22.0). To identify highly variable genes (HVGs), modelGeneVar (scran) was applied, and the top 2,000 HVGs were selected to compute principal components (PCs). The top 50 PCs were used for dimensional reduction using scater (runUMAP, v1.41.0) and for graph-based cell clustering using a Shared Nearest Neighbor (SNN) graph (buildSNNGraph, scran) in combination with the Louvain algorithm (cluster_louvain, igraph, v2.3.1). To assess cluster stability and robustness across parameter space, a parameter grid search was conducted incorporating (i) the number of HVGs: 1,000, 2,000, 3,000, 5,000, and 10,000; (ii) the number of PCs: 10, 20, 30, 50, and 100; (iii) the nearest neighbours (k): 10, 15, 20, and 30; and (iv) the resolution parameters: 0.3, 0.5, 0.7, 0.8, 1.0, 1.2, 1.5, 1.8, and 2.0. This yielded 900 distinct clustering schemes (5 * 5 * 4 * 9). Clustering consistency and consensus co-clustering frequencies were evaluated for every cell pair across all iterations (Fig S1D) and against the clustering scheme used in Fig 1B.

### Assignment of cluster names

Cluster names were assigned based on known cell group information within each cluster. For example, 3 from 4 plasma cells were in cluster 1, this cluster was named ‘Plasma’. For clusters lacking such experimental validation (clusters 3 and 5), cell types were inferred using SingleR (v2.14.0) ^47^ with the HumanPrimaryCellAtlas from celldex (v1.22.0) ^47^ as reference, and were assigned as Myeloid and Immu (immune), respectively. Cluster marker genes were identified using findMarkers (scran). Genes were defined as cluster-specific markers if they satisfied a threshold of p < 0.01 and log2 FC > 1 against all other clusters. Differential expression between M0-like and M1-like disseminated cancer cells (DCCs) was evaluated using the same framework with a more stringent cutoff of log_2 FC > 2.

### Copy number variance analysis

Single-cell copy-number variation profiles were inferred using inferCNA (v1.28.0; Bioconductor)[DOI: <u>10.18129/B9.bioc.inferCNA</u>]. Non-malignant control profiles—comprising healthy donor cells along with experimentally validated erythroid and plasma cell populations—served as the diploid reference baseline. Cell-level copy number variation (CNA) scores were quantified using the processed expression matrix output by inferCNA For each cell, the CNA score was computed as the average absolute deviation of all gene-level expression ratios from the diploid baseline (value of 1) as colMeans(abs(CNA_matrix - 1)). Higher CNA scores indicate higher degrees of chromosomal instability and copy number alterations. To classify cells harbouring copy-number variations (CNAs), individual cell CNA scores were evaluated against non-malignant reference cells. Threshold was established based on the baseline background signal of reference cells. Cells with a CNA score exceeding 0.05 were designated as CNA-positive according to Mean + 2SD(0.0478) of reference baseline (Fig S1F).

### Pathway enrichments of clusters

To infer the pathways enriched in each cluster, we performed hypergeometric tests on the cluster marker gene sets using the fora function from the fgsea package (version 1.38.0). Pathway enrichment was assessed against multiple reference databases, including GOMF, GOBP, GOCC, and HALLMARK gene sets downloaded via msigdbr (version 26.1.0), as well as two comprehensive cell-type marker databases, PanglaoDB ^48^ and CellMarker ^49^. For genes specifically enriched in the M0-DCC and M1-DCC clusters, we additionally examined their overlap with cancer metaprograms ^25^.

### Cancer type specific marker gene sets

For cancer type specific markers of cells that formed the M1-DCC cluster but originated from different tissues, findMarker was used. For this we had to exclude ESCA as it contributed too few cells from this cancer type (n=1). Marker genes were sorted according to p-value and the top 30 markers from each cancer type were plotted into the heatmap for cells from the M1-DCC and M0-DCC clusters. The same approach was applied to cells from the M0-DCC cluster comprising cells from four cancer types.

### Cancer type specific transcript factor activities for M0-DCC

To infer the transcription factor (TF) activities for cells of the M0-DCC cluster, we first downloaded TF network from Ominipath (https://omnipathdb.org/interactions?datasets=tf_target,dorothea). Then the Unweighted Linear Model (ULM) algorithm implemented in decoupleR (version 2.9.7) was applied based on a normalized expression matrix. TF activity scores were grouped by cancer type. One-way Analysis of Variance (ANOVA) was performed for each TF across cancer types, with raw p-values adjusted for TFs using the Benjamini-Hochberg False Discovery Rate. For TFs with p.adj< 0.05, Tukey’s Honestly Significant Difference (Tukey HSD) *post-hoc* tests were conducted to perform pairwise comparisons between distinct cancer types. Pairwise comparisons with adjusted p < 0.05 were considered cancer-type-specific TFs. We only took TFs that with targeted gene between 5 and 100.

### Target Gene Expression Profiling Across Single-Cell M0/M1 and TCGA Datasets

To evaluate the transcriptional output of M0-DCC cancer-type-specific transcription factors (TFs), we examined the expression patterns of their downstream target genes across single cell M0-DCC and M1-DCC clusters, as well as bulk RNA-sequencing profiles from corresponding TCGA cancer cohorts. For TCGA data analysis, gene expression matrices were normalized as log2(1+TPM). Patients were grouped by specific cancer types, with Lung Adenocarcinoma (LUAD) and Lung Squamous Cell Carcinoma (LUSC) integrated into a unified Non-Small Cell Lung Cancer (NSCLC) cohort.

### Stemness Indexing

Cellular stemness scores were quantified using three complementary pipelines applied to the normalized expression data: CytoTRACE (v0.1.0) ^50^, fitdevo (v1.2.0) ^51^, and Cell Biology Consortium stemness signature ^52^. Stemness scores were evaluated across cluster 1-6.

### A human cell type atlas as reference panel

To construct a comprehensive single cell reference panel, three publicly available single-cell atlases were integrated, comprising (i) the Human Bone Marrow Atlas ^53^ containing single-cell profiles derived exclusively from adult bone marrow tissue; (ii) the Human Lymph Node Atlas ^54^, comprising single-cell transcriptomes from adult lymph node tissue; (iii) the Human Cell Landscape (HCL) ^40^, a comprehensive multi-tissue atlas spanning over 60 distinct tissue types, incorporating cells across adult and fetal tissues as well as human Embryonic Stem Cells (hESC). To accurately assess the developmental stage, cell types were classified as stage-specific if detected in > 1,000 cells in one stage but < 100 cells in the comparative stage. Cell types meeting representation thresholds in both stages were classified as stage-shared. Given the unbalanced cell counts across three atlas datasets and the low number of our RNA-seq data (N=864 across 6 clusters) from EPCAM+ cells, we implemented a stratified downsampling strategy. Each annotated cell type within each reference atlas was randomly down sampled to a maximum of 1,000 cells per developmental stage. Stemness scores for individual cells within the single-cell atlas were computed following the same pipeline applied to the EPCAM+ RNA-seq dataset in this study. To derive the reference cell-type signatures from the RNA-seq data of EPCAM+, gene set enrichment scores were calculated using the AddModuleScore_UCell function from the UCell R package (v2.16.0) against the marker gene set of each cell type derived from this paper.

### Integration of EpCAM+ cells into the human cell atlas

Each batch was first normalized with SCTrasform from R package Seurat (version 5.5.0) and then integrated using RunHarmony from R package harmony (version 2.0.3) by using the top 50 harmony components to get UMAP dimension reduction. After integration, cells were clustered using FindNeighbor and FindClusters from Seurat using the top 50 harmony components and parameter resolution equals 0.1. For cell integration with the bone marrow or lymph node atlas each cluster was assigned according to the most abundant cell type as per atlas annotation or from this work. For fine mapping of M0-DCC integration among fetal tissue-derived cells (i.e. the cluster comprising most M0-DCC), RunUMAP was re-done using top 50 harmony components. Cell types with less than 50 cells were merged to rare cell types and the cell type annotation was marked at the median position of cells of this type in the UMAP.

### Similarity of M0- and M1-DCC to human and murine embryonic cells

Gene expression data of early human embryos were obtained from two data sets, (i) from zygote to morula stage ^28^ and (ii) from 8-cell embryos to hypoblasts ^27^. Single cell RNA-seq count tables were download and expression data normalized using Seurat’s NormalizeData (version 5.5.0). Signature scores were calculated at single-cell resolution using the AddModuleScore_UCell function from the UCell R package (v2.16.0). Signature scores were evaluated for marker gene sets in this paper. Additionally, signature scores for Erythroblast and Plasma cells were calculated using the top 500 marker genes given by human bone marrow atlas ^55^. The gene expression was also visualized in a heatmap. Since much higher early embryonic cell numbers are available for murine embryos, pathway enrichment in Fig S3E was assessed by using data from reference ^56^ for zygote to 16-cell embryos.

### Induction of the M0-DCC phenotype in vitro

MDA-MB-231, MDA-MB-231-1833, CAL-51, and SkBr3 cells were cultured in DMEM medium (Pan-Biotech, Germany) supplemented with 5% FCS (Sigma-Aldrich, Germany), 2 mM l-glutamine (Pan-Biotech, Germany), and 1% penicillin/streptomycin (Pan-Biotech, Germany). MCF7, MCF7-GFP, BT474, Du4475, and T47D cells were propagated in RPMI-1640 medium (Pan-Biotech, Germany) supplemented with 10% FCS, 2 mM l-glutamine, and 1% penicillin/streptomycin. All cell lines were kept at 37 °C and 5% CO2 in a fully humidified incubator and negatively tested for mycoplasma by PCR.

For the M0-DCC-phenotype induction experiments, the cells were seeded at densities of 30000, 5000, and 3000 cells/cm^2^ and were plated in 96 well plates, 24 well plates, or 6 well plates to account for high, medium and low densities, respectively. The cytokines were added from day 1 following plating and were topped off every 2 days. The final concentration of ANGPTL7 (Preprotech Cat# 130-22), IL33 (Preprotech Cat# 200-33) and SERPINB2 also known as PAI 2 (Preprotech Cat# 140-06) were 75 ng/ml whereas that of HIL6 (a kind gift of S. Rose-John, Christian-Albrechts-University, Germany) (Fischer (1997), Nat Biotechnol. 15(2):142-5) was 40 ng/mL. Differentiating cells were harvested at day 10 and subjected to downstream analyses.

To test the reversibility of the induced phenotype, MCF7 and SkBr3 cells seeded at LD at were treated as previously described for 8 days. Control and treated cells were stained for anti-human cKIT-BV421 (BioLegend cat# 313216 clone: 104D2), anti-human CD36-PE (BioLegend cat# 336206 clone: 5-271), and anti-human EpCAM-AF488 (BioLegend cat# 324210 clone: 9C4). Marker expression was determined using flow cytometry. Subsequently, both treated and untreated cell groups were maintained without treatment for an additional eight days, with medium changes every other day. After this withdrawal period, cells were stained again, and c-KIT, CD36, and EpCAM expression was re-evaluated by flow cytometry. Marker expression levels obtained immediately after treatment were then compared to those measured following the eight-day withdrawal period.

### Gene expression analysis by qRT-PCR

Primers were designed with the help of the NCBI PrimerBlast online tool targeting the canonical transcript isoform in the respective genes of interest. Primer validation consisted of three steps (i) gradient PCR to identify the optimal annealing temperature, (ii) standard curve assessment to determine the dynamic range, the sensitivity and the efficiency of each qPCR assay, and (iii) restriction fragment length polymorphism (RFLP) assessment of PCR products generated by each assay to confirm the specificity of the amplified fragments. Total RNA extraction was carried out using the NucleoSpin RNA Mini Kit (Macherey-Nagel, Germany). Subsequently, cDNA was generated from a maximum 500 ng of total RNA using the SuperScript™ First-Strand Synthesis System for RT-PCR (Thermo-Fischer Scientific). Resulting cDNA samples were diluted 1:5 in wate and gene expression was subsequently quantified by qPCR (Biorad CFX Opus 96 Real-Time PCR System). Relative gene expression was quantified using the comparative ΔΔCP method. All measurements were conducted in triplicates. Average CP values of target genes were normalized to equivalent values of the respective reference genes (*GAPDH* and *ACTB*) to obtain ΔCP values (ΔCP = CP_target − CP_reference). Relative expression was then calculated against a calibrator sample using ΔΔCP (ΔΔCP = ΔCP_sample − ΔCP_calibrator) and expressed as 2^−ΔΔCP fold change.

### Flow cytometry

Adherent cells were trypsinized with trypsin/EDTA (Pan-Biotech, Germany) for 3 min. Viability dye eFluor 780 (ebioscience, Germany) was used to identify dead cells at a concentration of 1:2000 as recommended by the manufacturer. The cell suspension was then washed with flow cytometry buffer composed of PBS/2% FCS/0.01% NaN3. To reduce nonspecific binding, single-cell suspensions were incubated for 10 min at RT with Human TruStain FcX™ (BioLegend, Germany). Cells were stained using the following antibodies anti-human CD36-PE, anti-human EpCAM-AF488, and anti-human cKIT-BV421. To control the staining, the following isotypes were used: Mouse IgG1, κ Isotype Ctrl Antibody-BV421, and Mouse IgG2a, κ Isotype Ctrl Antibody-PE (BioLegend, Germany) for 20 min at RT. Cells were then fixed for 20 min using the FluoroFix™ Buffer (BioLegend, Germany) and then washed a final time with flow cytometry buffer. Stained cells were measured using the Beckman Coulter Gallios Flow Cytometer and the data was analyzed with FlowJo 10.5.3 (Treestar, USA).

### MCF7 co-culture with bone marrow

2000 MCF7 cells in Human Plasma-Like Medium (HPLM) supplemented with 10% AB serum, 0.5% penicillin/streptomycin, and 1X 2-HBA salt were seeded per well into a 96 well plate. The following day, frozen bone marrow previously obtained from hip replacement surgery at the University Center of orthopedics in Bad Abbach was thawed and diluted in 10 mL of supplemented HPLM and was washed and counted. 80,000 cells of bone marrow suspension cells were then added to the MCF7 and were cultured at 37 °C and 5% CO2 in a fully humidified incubator and negatively tested for mycoplasma by PCR. The MCF7-BM co-culture was kept for 5 days.

### Cell isolation for sequencing with 10X Genomics

Following treatment, MCF7 cells were harvested using trypsin/EDTA for 3 min. The cell suspension was washed with 2 mL of flow cytometry buffer (PBS supplemented with 20% FCS). To reduce nonspecific antibody binding, single-cell suspensions were incubated for 10 min at room temperature with Human TruStain FcX™ blocking solution (2.5 µL FcX™ reagent diluted in 7.5 µL flow cytometry buffer per sample). Cells were subsequently stained with TotalSeq™-B oligonucleotide-conjugated antibodies against TotalSeq™-B0407 anti-human CD36 (BioLegend cat# 336233 clone: 5-271) and TotalSeq™-B0061 anti-human CD117 (c-kit) (BioLegend cat# 313247 clone: 104D2) for 30 min at 4°C. Following staining, cells were washed three times with PBS containing 20% FCS and stained with 7-aminoactinomycin D (7-AAD) (BioLegend cat# 420403) immediately prior to fluorescence-activated cell sorting (FACS). Viable (7-AAD-negative) cells were sorted using a BD FACSAria™ Fusion High-Speed Cell Sorter, and 100,000 viable cells per condition were collected and resuspended in 1 mL PBS containing 20% FCS for single-cell capture. For co-culture experiments, GFP-labelled MCF7 cells were used, and GFP-positive viable cells were sorted and collected in 1 mL PBS containing 20% FCS. Single-cell gene expression and antibody-derived tag (ADT) libraries were generated using the Chromium Next GEM Single Cell 3ʹ Reagent Kit v3.1 (10x Genomics) in combination with the Chromium Single Cell 3ʹ Feature Barcode Library Kit, according to the manufacturer’s instructions. Gene expression and ADT libraries were prepared and sequenced separately, enabling simultaneous quantification of transcriptomes and cell surface protein expression.

### 10X#Genomics RNA-seq data analysis

We used cellRanger-10.0.0 to get 10x RNA counts and ADT count. Cells with detected gene numbers of <500, reads <1000 or mitochodrila> 10% were filtered. Doublets were detected using scDblFinder (v1.26.1) and removed. RNA gene expression data were normalized using Seurat’s `NormalizeData` (version 5.5.0), followed by highly variable feature selection via `FindVariableFeatures`. The expression matrix was then scaled and centered using `ScaleData`. Principal Component Analysis

(PCA) was conducted with `RunPCA`, and the first 30 principal components were used to compute a Uniform Manifold Approximation and Projection (UMAP) embedding via `RunUMAP` for visualization. For samples with antibody-derived tag (ADT) data, surface protein expression levels were normalized using Seurat’s `NormalizeData` function with the Center Log Ratio (`CLR`) transformation method across cells. To identify cells with high expression surface markers, expression thresholds were set at the 70th percentile across cells. Cells with antibody-derived tag (ADT) expression above these cutoffs were classified as marker-high for downstream analysis.

### Gene Set Enrichment analysis for double high vs double low genes

The position density of CD36/KIT double high cells and CD36/KIT double low cells was assessed for each sample using Plot_Density_Custom from scCustomize(version 3.3.0). Differential gene expression analysis was performed for each individual sample using the `FindMarkers` function in Seurat (comparing double-high versus double-low cell populations) with parameter only.pos = F, min.pct = 0, logfc.threshold = 0. To prepare input ranks for pathway analysis, genes were ordered by a signed significance score:

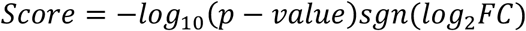

where

*sgn*(*log*_2_*FC*) represents the sign (+1 or −1) of *log*_2_*FC*. Genes were subsequently ordered in descending rank by this score for GSEA execution, using command fgsea from fgsea package (1.38.0) against M0/M1 signatures.

### Analysis of Cancer Cell Line Encyclopedia (CCLE) cells for metastatic potential

The phenotype and expected count matrix of CCLE data are download from CCLE release version 25Q3. Gene expression data were normalized using the Trimmed Mean of M-values (TMM) method implemented in the edgeR(‘4.10.1’) package ^56^, and then log2-transformed with a prior count of 1 [log2(CPM + 1)]. Z-score was calculated for M0/M1 signatures using function z-score from GSVA(‘2.6.2’). The metastatic potential ^57^ accross varies tissues was compared between M0 z-score positive vs negative groups and M1 z-score positive vs negative using the non-parametric Wilcoxon rank-sum test. A similar comparison was performed for cell lines generated from primary tumors versus metastasis-derived cell lines ^57^.

### Analysis of patient outcome

For all survival analysis, only M0-stage patients were included, i.e. all patients without evidence of manifest metastasis at bone marrow or lymph node sampling. Survival outcomes were evaluated using Kaplan–Meier analysis. Survival curves were generated using the survfit function from the survival (3.8.9) package and visualized using ggsurvplot from the survminer(0.5.2). Statistical differences between groups were assessed using log-rank test. For NSCLC patient with KIT protein staining Univariate Cox proportional hazards regression models were constructed to evaluate the prognostic association between individual clinicopathological and immunofluorescence features and tumor specific survival (TSS). The analyzed feature set included protein marker detection (KIT, EpCAM, and CK), conventional clinical information (age, sex, TNM T grade and N stage). Hazard ratios (HRs), 95% confidence intervals (CIs), and corresponding p-values were calculated using the coxph function from the survival (3.8.9) package. Multivariate Cox proportional hazards regression modelling including KIT, T grad, N stage, age and sex was performed to identify predictors of tumour-specific survival (TSS) by using the coxph function from the survival package. The result was visualized using ggforest function from survminer.

### Statistics and reproducibility

No statistical method was used to predetermine the sample size. For RNA-seq experiments, sample size was determined by the availability of high-quality RNA from EPCAM+ cells and control cells to ensure sufficient power for meaningful patterns. (For other experiments, sample numbers were based on practical considerations to reliably address primary objectives. Post hoc assessments confirmed that sample numbers were adequate for valid results.)

To ensure reproducibility, independent replicate experiments were performed, including biological and technical replicates where applicable. Technical replicates were used to assess measurement accuracy, assay reproducibility and technical variability. Data were analysed using multiple statistical approaches to confirm robustness. All replication attempts were successful, with consistent results across biological samples and experimental conditions. WB analyses were performed in at least two independent biological replicates, and microscopy images were obtained from multiple replicates, with consistent findings observed.

Statistical analyses were conducted using GraphPad Prism (version 9.3.1) and R (version 4.6.1). Univariable, multivariable and survival analyses were performed using Cox regression and the Mantel– Cox log-rank test. All tests were two-sided, with *P* < 0.05 considered statistically significant.

Patient samples were included in the study according to availability, which was determined by factors such as RNA quality and available survival data. No preassigned groupings or patient selections were made before data analysis. (As the study did not involve controlled experimental conditions or treatment interventions, random allocation of samples was not applicable. No formal covariate-based randomization was performed, as the study addressed the natural variability within the available patient samples). All patient samples were pseudonymized according to EU General Data Protection Regulation (GDPR) and pseudonyms linked clinical and outcome data. Investigators were blinded to patient disease progression and clinical status until final bioinformatics analysis. This ensured that the data collection and initial analysis were conducted without bias related to disease state. For post-RNA-seq analyses, such as survival analysis, reidentification was necessary to link clinical outcomes to the molecular data. (Blinding was not feasible for survival analysis, as patient outcomes were required to interpret these findings; however, bias was minimized by the use of pseudonymized data during the initial steps of the experiment).

## Legends for Supplementary Figures

**Figure S1: Assignment of cell identities to the gene expression clusters of EpCAM^+^ cells.**

**a) Impact of tissue origin.** Dimensionality reduction plot of EpCAM+ cells based on their transcriptional profiles and highlighting cluster positioning by tissue origin.

**b) Distribution of cells across tissue origin and cluster identity**. Only samples from M0-stage patients were considered (LN = lymph node, BM = bone marrow). Each entry represents the proportion of cells assigned to each cluster for BM and LN samples. Note, the scarcity of M1-DCC cluster assignments for cell from BM as opposed to LN.

**c) Cell type assignment for BM-derived myeloid cells and LN-derived immune cells.** Cell type inference of cluster 3 (myeloid cells) and 5 (diverse immune cells) by SingleR using HumanPrimaryCellAtlas as cell type reference.

**d) Assessment of cluster stability.** Heatmap displaying the co-clustering frequency for each pair of samples across 900 parameter grids. Values range from 0 (never clustered together) to 1 (always clustered together). Annotations indicate cell identity (cluster assignment), group information (M0 patient, M1 patient, non-cancer donor (HD), or experimentally validated ERP/Plasma cells), CNA scores, and CNA detection status (detected/not detected). The high co-clustering frequencies within cluster assignment confirm the robustness of the clustering results.

**e) Heatmap of all samples using the most informative genes.** Heatmap displaying the expression of marker genes across individual cells. Rows represent marker genes, with row annotations indicating the cluster/cell type to which each marker gene belongs. Columns represent individual cells, with column annotations indicating the cluster identity of each cell.

**f) Density plot showing the distribution of CNA scores for reference cells and other EPCAM+ cells.** The blue vertical line indicates the upper bound of the one-sided 95% confidence interval of the reference cell CNA scores. Round of the upper bound (0.05) was used as the cutoff to classify cells with significantly elevated CNA burden. Cells with CNA scores above this threshold are considered CNA-positive, while those below are classified as CNA-negative.

**g) Cluster composition after projection of EpCAM+ cells into the BM and LN-atlas.** Left panel: UMAP projection of integrated single-cell transcriptomic data combining EPCAM+ cells from this study with BM and LN reference atlases. Each cluster was assigned a **cell type label** based on the **majority annotation** among cells within that cluster, regardless of whether the annotation originated from the reference atlases (BM/LN) or from our EPCAM+ cluster assignments. The clustering shows that M0-DCC form an independent cluster that does not overlap with atlas while other EPCAM+ cells integrate into BM or LN atlases. Right panel: Bar plot showing the cell type composition of clusters that contain most M0-DCC, M1-DCC, Myeloid, ERP, Plasma, and immune cells following integration of EPCAM+ cells from this study with BM and LN reference atlases. Each bar represents a cluster, with colors indicating the proportion of different cell types present within that cluster. Despite the large size of the reference atlases, only a small number of atlas-derived cells are found within M0-DCC cluster, i.e. M0-DCC form a cluster unknown to the BM or LN atlas.

**Figure S2: Pathway enrichment in EpCAM^+^ cell clusters from BM and LN.**

a)-d) Dot plots showing the top enriched pathways identified in the ERP/Plasma/Myeloid/Immu clusters separately, based on the databases GOBP, GOMF, GOCC, HALLMARK and the cell type marker database panglao and cellMarker. X-axis shows the −log10(p value) and colors indicate fold-enrichment.

a) Histogram plot showing the top enriched pathways of M1-DCC vs M0-DCC clusters based on top 10 cancer specific metaprogram. X-axis shows the −log10(p value). M1-DCC are enriched for genes in cancer stress/emt/epithelial senescence metaprograms, while M0-DCC in genes characteristic for rbcs/myc/cell cycle metaprograms.

**Figure S3: Enrichment of embryonic features in M0-DCC**

**a) Stemness scores by cell type and algorithm.** Heatmap displaying the stemness scores for HCL atlas cells and UCell scores for EPCAM+ cluster signatures averaged across different cell types. Annotations indicate: (1) whether the cell type is stage-specific, i.e. found in adult, fetal, both or HESC; (2) cell numbers found among fetal, or (3) adult cells from atlas. Cell type contain more than 1000 cell in each stage will be counted as 1000. Note that erythroid, erythroid progenitor cell, fetus specific cell types (e.g. primordial germ cells, fetal adrenal gland inflammatory cell, fetal proliferating T cells) and human embryonic cells (HESC) are high in stemness and M0-DCC scores.

**b) UMAP projection of integrated single-cell data highlighting stemness and CNA scores.** EPCAM^+^ cells from this study were combined with HCL (Human Cell Landscape) and BM (bone marrow) reference atlases. Top: HCL cells are colored by fitdevo stemness scores, with brighter color indicating greater stemness scores. High stemness scores are concentrated in regions occupied by fetal cells and erythroid cell types from HCL atlas (see figure 4b for cell type information). Bottom: EPCAM+ cells are colored by CNA scores. M1-DCC cells, distributed across diverse regions of the adult HCL, show high CNA scores. M0-DCC cells, localized predominantly within the fetal HCL area, also exhibit high CNA scores.

**c) Heat map of marker genes of EpCAM^+^ ERP and plasma cells in human embryonic cells**. Embryonic cells widely express ERP marker genes of the Bone Marrow Atlas ^53^, in contrast to plasma cell markers. Left panel: cells from REF ^28^, right panel: cells from REF ^27^.

**d) Early human embryonic cells express more marker genes of M0-DCC than of M1-DCC.** Top panel: cells from REF ^28^, right panel: cells from REF ^27^.

**e) Enrichment of M0-DCC and M1-DCC genes in early embryos.** Genes upregulated in M0-DCC are highly enriched in embryos from zygote to 16-cell stage as opposed to characteristic M1-DCC genes. Embryonic data from REF ^55^. Bar plots showing the −log10(p-value) for overlap between M0-DCC (left) and M1-DCC (right) marker gene sets and embryonic cell marker genes spanning developmental stages from zygote to 16-cell. The M0-DCC marker set shows significantly higher enrichment across all embryonic stages compared to M1-DCC, indicating a much stronger transcriptional resemblance of M0-DCC to early embryonic programs.

**f) Expression of characteristic gene signatures of EpCAM+ cell clusters by early embryos.** Box plots showing the distribution of UCell scores for all EPCAM+ cluster marker gene sets in human embryonic cell types spanning developmental stages from oocyte to hypoblast using cells from REF ^28^ and REF ^27^. The box represents the IQR, the center line denotes the median, and whiskers extend to 1.5 × IQR early embryonic genes in M0-DCC and M1 DCC.

**Figure S4: Embryonic and erythroid markers enable monitoring of M0-DCC induction in cell lines**

**a) M0-DCCness vs M1-DCCness in human cell lines.** Bar plot showing the **z-scores** of **M0-DCC** and **M1-DCC** marker gene sets across cancer cell lines (SKBR3, BT474, T47D, MCF7, MDAMB231, CAL51, DU4475).

**b) Expression of marker genes in EpCAM+ cell clusters and early human embryos.** Top: Dot plot of selected genes in EPCAM+ cell clusters. Bottom: Expression of the same marker genes in human embryonic cells, using cells from REF ^27^, and cells from REF ^26^.

**c) and d) Induction of marker genes in human breast cancer cell lines.** Different cytokine combinations induce marker genes in various cell lines to various degrees (c). Marker genes are per se elevated in bone marrow-selected MDA-MB-231-1833 cells compared to parental MDA-MB-231 (d).

**d) Induction of CD36 and KIT protein in CAL51 and Du4475.** Cells with low M1-DCC score cannot be induced to express the M0-DCC marker proteins CD36 and KIT.

**e) CD36 and KIT protein induction in MCF7 cells by ADT sequencing.** Box plot showing the distribution of the antibody DNA tags ADT_CD36 and ADT_CD117 across four experimental conditions: HD-Treated, HD-Untreated, LD-Treated, and LD-Untreated. Each box represents the interquartile range (IQR) with the median indicated by the center line, and whiskers extend to 1.5 × IQR. Kruskal-Wallis test (p < 2.2e-16) indicate significant differences across conditions. LD-Treated condition show the highest ADT_CD36 and ATD_CD117 expression

## Notes

### Competing Interest Statement

H. Koerkel-Qu, E. Raya, M. Werner-Klein, T. Mederer, D. Spitzl, M. Guzvic, C. A. Klein are inventors on IP applications of the Fraunhofer Society.

## References

1 Mai, N., Fernandez, N., Drilon, A. & Chakravarty, D. Precision Oncology: 2025 in Review. Cancer Discov 15, 2414–2421 (2025). 10.1158/2159-8290.CD-25-1784

2 Schlimok, G. et al. Micrometastatic cancer cells in bone marrow: in vitro detection with anti-cytokeratin and in vivo labeling with anti-17-1A monoclonal antibodies. Proc Natl Acad Sci U S A 84, 8672–8676 (1987).

3 Klein, C. A. et al. Comparative genomic hybridization, loss of heterozygosity, and DNA sequence analysis of single cells. Proc Natl Acad Sci U S A 96, 4494–4499 (1999).

4 Klein, C. A. et al. Combined transcriptome and genome analysis of single micrometastatic cells. Nat Biotechnol 20, 387–392 (2002).

5 Guetter, S. et al. MCSP(+) metastasis founder cells activate immunosuppression early in human melanoma metastatic colonization. Nat Cancer 6, 1017–1034 (2025). 10.1038/s43018-025-00963-w

6 Guzvic, M. et al. Combined genome and transcriptome analysis of single disseminated cancer cells from bone marrow of prostate cancer patients reveals unexpected transcriptomes. Cancer research 74, 7383–7394 (2014). 10.1158/0008-5472.CAN-14-0934

7 Hosseini, H. et al. Early dissemination seeds metastasis in breast cancer. Nature 540, 552–558 (2016). 10.1038/nature20785

8 Husemann, Y. et al. Systemic spread is an early step in breast cancer. Cancer cell 13, 58–68 (2008).

9 Werner-Klein, M. et al. Interleukin-6 trans-signaling is a candidate mechanism to drive progression of human DCCs during clinical latency. Nat Commun 11, 4977 (2020). 10.1038/s41467-020-18701-4

10 Werner-Klein, M. et al. Genetic alterations driving metastatic colony formation are acquired outside of the primary tumour in melanoma. Nat Commun 9, 595 (2018). 10.1038/s41467-017-02674-y

11 Riethdorf, S., Wikman, H. & Pantel, K. Review: Biological relevance of disseminated tumor cells in cancer patients. International journal of cancer 123, 1991–2006 (2008).

12 Ulmer, A. et al. Quantitative measurement of melanoma spread in sentinel lymph nodes and survival. PLoS medicine 11, e1001604 (2014). 10.1371/journal.pmed.1001604

13 Hartkopf, A. D. et al. Disseminated tumour cells from the bone marrow of early breast cancer patients: Results from an international pooled analysis. Eur J Cancer 154, 128–137 (2021). 10.1016/j.ejca.2021.06.028

14 Klein, C. A. The systemic progression of human cancer: a focus on the individual disseminated cancer cell--the unit of selection. Adv Cancer Res 89, 35–67 (2003).

15 Lammers, R. et al. Monoclonal antibody 9C4 recognizes epithelial cellular adhesion molecule, a cell surface antigen expressed in early steps of erythropoiesis. Exp Hematol 30, 537–545 (2002). 10.1016/s0301-472x(02)00798-1

16 Gires, O., Pan, M., Schinke, H., Canis, M. & Baeuerle, P. A. Expression and function of epithelial cell adhesion molecule EpCAM: where are we after 40 years? Cancer Metastasis Rev 39, 969–987 (2020). 10.1007/s10555-020-09898-3

17 Gottlinger, H. G., Funke, I., Johnson, J. P., Gokel, J. M. & Riethmuller, G. The epithelial cell surface antigen 17-1A, a target for antibody-mediated tumor therapy: its biochemical nature, tissue distribution and recognition by different monoclonal antibodies. International journal of cancer 38, 47–53 (1986). 10.1002/ijc.2910380109

18 Kwon, D. I. et al. Homeostatic serum IgE is secreted by plasma cells in the thymus and enhances mast cell survival. Nat Commun 13, 1418 (2022). 10.1038/s41467-022-29032-x

19 Hao, Y. et al. Integrated analysis of multimodal single-cell data. Cell 184, 3573–3587 e3529 (2021). 10.1016/j.cell.2021.04.048

20 Schardt, J. A. et al. Genomic analysis of single cytokeratin-positive cells from bone marrow reveals early mutational events in breast cancer. Cancer cell 8, 227–239 (2005).

21 Schmidt-Kittler, O. et al. From latent disseminated cells to overt metastasis: genetic analysis of systemic breast cancer progression. Proc Natl Acad Sci U S A 100, 7737–7742 (2003).

22 Stoecklein, N. H. et al. Direct genetic analysis of single disseminated cancer cells for prediction of outcome and therapy selection in esophageal cancer. Cancer cell 13, 441–453 (2008).

23 Weckermann, D. et al. Perioperative activation of disseminated tumor cells in bone marrow of patients with prostate cancer. J Clin Oncol 27, 1549–1556 (2009).

24 Klein, C. A. et al. Genetic heterogeneity of single disseminated tumour cells in minimal residual cancer. Lancet 360, 683–689 (2002).

25 Gavish, A. et al. Hallmarks of transcriptional intratumour heterogeneity across a thousand tumours. Nature 618, 598–606 (2023). 10.1038/s41586-023-06130-4

26 Zhang, C. X., Huang, R. Y., Sheng, G. & Thiery, J. P. Epithelial-mesenchymal transition. Cell 188, 5436–5486 (2025). 10.1016/j.cell.2025.08.033

27 Petropoulos, S. et al. Single-Cell RNA-Seq Reveals Lineage and X Chromosome Dynamics in Human Preimplantation Embryos. Cell 165, 1012–1026 (2016). 10.1016/j.cell.2016.03.023

28 Yan, L. et al. Single-cell RNA-Seq profiling of human preimplantation embryos and embryonic stem cells. Nat Struct Mol Biol 20, 1131–1139 (2013). 10.1038/nsmb.2660

29 Thowfeequ, S., Hanna, C. W. & Srinivas, S. Origin, fate and function of extraembryonic tissues during mammalian development. Nat Rev Mol Cell Biol 26, 255–275 (2025). 10.1038/s41580-024-00809-w

30 Klein, C. A. Parallel progression of primary tumours and metastases. Nat Rev Cancer 9, 302–312 (2009).

31 Klein, C. A. Cancer progression and the invisible phase of metastatic colonization. Nat Rev Cancer 20, 681–694 (2020). 10.1038/s41568-020-00300-6

32 Scheunemann, P., Izbicki, J. R. & Pantel, K. Tumorigenic potential of apparently tumor-free lymph nodes. N Engl J Med 340, 1687 (1999). 10.1056/nejm199905273402116

33 Solakoglu, O. et al. Heterogeneous proliferative potential of occult metastatic cells in bone marrow of patients with solid epithelial tumors. Proc Natl Acad Sci U S A 99, 2246–2251 (2002). 10.1073/pnas.042372199

34 Palis, J. Primitive and definitive erythropoiesis in mammals. Front Physiol 5, 3 (2014). 10.3389/fphys.2014.00003

35 Seu, K. G. et al. Unraveling Macrophage Heterogeneity in Erythroblastic Islands. Front Immunol 8, 1140 (2017). 10.3389/fimmu.2017.01140

36 Elgammal, Y. et al. Characterization of the human erythromyeloblastic islands (EMBIs). Blood 146, 182–182 (2025). 10.1182/blood-2025-182

37 Lopez-Yrigoyen, M. et al. Genetic programming of macrophages generates an in vitro model for the human erythroid island niche. Nat Commun 10, 881 (2019). 10.1038/s41467-019-08705-0

38 Fischer, M. et al. I. A bioactive designer cytokine for human hematopoietic progenitor cell expansion. Nat Biotechnol 15, 142–145 (1997). 10.1038/nbt0297-142

39 Barretina, J. et al. The Cancer Cell Line Encyclopedia enables predictive modelling of anticancer drug sensitivity. Nature 483, 603–607 (2012). 10.1038/nature11003

40 Han, X. et al. Construction of a human cell landscape at single-cell level. Nature 581, 303–309 (2020). 10.1038/s41586-020-2157-4

41 Brabletz, T. et al. Variable beta-catenin expression in colorectal cancers indicates tumor progression driven by the tumor environment. Proc Natl Acad Sci U S A 98, 10356–10361 (2001). 10.1073/pnas.171610498

42 Garber, J. E. & Offit, K. Hereditary cancer predisposition syndromes. J Clin Oncol 23, 276–292 (2005). 10.1200/JCO.2005.10.042

43 Asami, M. et al. A program of successive gene expression in mouse one-cell embryos. Cell Rep 42, 112023 (2023). 10.1016/j.celrep.2023.112023

44 Asami, M. et al. Human embryonic genome activation initiates at the one-cell stage. Cell Stem Cell 29, 209–216 e204 (2022). 10.1016/j.stem.2021.11.012

45 Perry, A. C. F., Asami, M., Lam, B. Y. H. & Yeo, G. S. H. The initiation of mammalian embryonic transcription: to begin at the beginning. Trends Cell Biol 33, 365–373 (2023). 10.1016/j.tcb.2022.08.008

46 Hartmann, C. H. & Klein, C. A. Gene expression profiling of single cells on large-scale oligonucleotide arrays. Nucleic Acids Res 34, e143 (2006).

47 Aran, D. et al. Reference-based analysis of lung single-cell sequencing reveals a transitional profibrotic macrophage. Nat Immunol 20, 163–172 (2019). 10.1038/s41590-018-0276-y

48 Franzen, O., Gan, L. M. & Bjorkegren, J. L. M. PanglaoDB: a web server for exploration of mouse and human single-cell RNA sequencing data. Database (Oxford) 2019 (2019). 10.1093/database/baz046

49 Zhang, X. et al. CellMarker: a manually curated resource of cell markers in human and mouse. Nucleic Acids Res 47, D721–D728 (2019). 10.1093/nar/gky900

50 Gulati, G. S. et al. Single-cell transcriptional diversity is a hallmark of developmental potential. Science (New York, N.Y 367, 405–411 (2020). 10.1126/science.aax0249

51 Zhang, F. et al. FitDevo: accurate inference of single-cell developmental potential using sample-specific gene weight. Brief Bioinform 23 (2022). 10.1093/bib/bbac293

52 Malta, T. M. et al. Machine Learning Identifies Stemness Features Associated with Oncogenic Dedifferentiation. Cell 173, 338–354 e315 (2018). 10.1016/j.cell.2018.03.034

53 Hay, S. B., Ferchen, K., Chetal, K., Grimes, H. L. & Salomonis, N. The Human Cell Atlas bone marrow single-cell interactive web portal. Exp Hematol 68, 51–61 (2018). 10.1016/j.exphem.2018.09.004

54 Tabula Sapiens, C., et al. The Tabula Sapiens: A multiple-organ, single-cell transcriptomic atlas of humans. Science (New York, N.Y 376, eabl4896 (2022). 10.1126/science.abl4896

55 Posfai, E. et al. Evaluating totipotency using criteria of increasing stringency. Nature cell biology 23, 49–60 (2021). 10.1038/s41556-020-00609-2

56 Robinson, M. D., McCarthy, D. J. & Smyth, G. K. edgeR: a Bioconductor package for differential expression analysis of digital gene expression data. Bioinformatics 26, 139–140 (2010). 10.1093/bioinformatics/btp616

57 Jin, X. et al. A metastasis map of human cancer cell lines. Nature 588, 331–336 (2020). 10.1038/s41586-020-2969-2

