## Supplementary Table 1 for "Metastatic founder cell candidates resemble preimplantation embryonic blastomeres"

| Cancer Type | Characteristic | Non-Metastatic (M0) | Metastatic (M1) |
|---|---|---|---|
| **Breast Cancer Patients** | **Recruited** | **449 (100%)** | **23 (100%)** |
| | Screened | 381 (85%) | 20 (87%) |
| | Patients with EpCAM+ cells detected | 158 (35%) | 14 (60%) |
| | Patients with EpCAM+ cells included in study | 86 (19%) | 10 (43%) |
| **EpCAM+ Single cells** | **Total cells isolated** | **593 (100%)** | **87 (100%)** |
| | **Cells included in study** | **269 (45%)** | **26 (30%)** |
| **NSCLC Patients** | **Recruited** | **133 (100%)** | **5 (100%)** |
| | Screened | 133 (100%) | 5 (100%) |
| | Patients with EpCAM+ cells detected | 59 (44%) | 4 (80%) |
| | Patients with EpCAM+ cells included in study | 28 (21%) | 2 (40%) |
| **EpCAM+ Single cells** | **Cells isolated** | **180 (100%)** | **22 (100%)** |
| | **Cells included in study** | **40 (22%)** | **10 (45%)** |
| **Prostate Cancer Patients** | **Recruited** | **487 (100%)** | **2 (100%)** |
| | Screened | 347 (71%) | 2 (100%) |
| | Patients with EpCAM+ cells detected | 177 (36%) | 2 (100%) |
| | Patients with EpCAM+ cells included in study | 111 (23%) | 2 (100%) |
| **EpCAM+ Single cells** | **Cells isolated** | **521 (100%)** | **16 (100%)** |
| | **Cells included in study** | **248 (48%)** | **9 (56%)** |
| **Esophageal Cancer Patients** | **Recruited** | **65 (100%)** | **5 (100%)** |
| | Screened | 54 (83%) | 5 (100%) |
| | Patients with EpCAM+ cells detected | 29 (44.6%) | 3 (60%) |
| | Patients with EpCAM+ cells included in study | 28 (43.1%) | 3 (60%) |
| **EpCAM+ Single cells** | **Cells isolated** | **94 (100%)** | **15 (100%)** |
| | **Cells included in study** | **48 (51.1%)** | **6 (40%)** |
