## Supplementary figures and images for "Metastatic founder cell candidates resemble preimplantation embryonic blastomeres"

### Supplementary Figure S1

#### Supplementary Figure 1

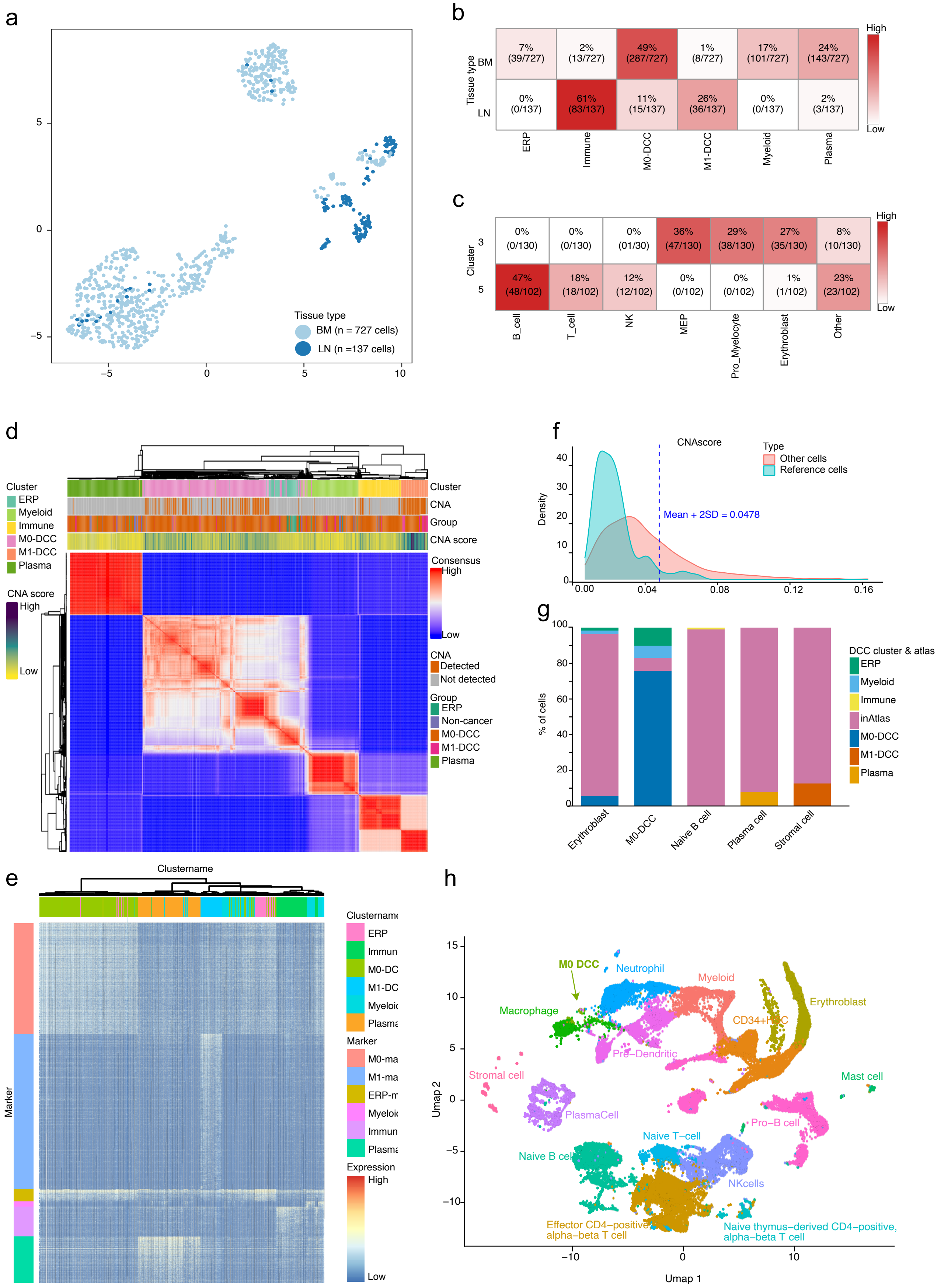

### Supplementary Figure S2

#### Supplementary Figure 2

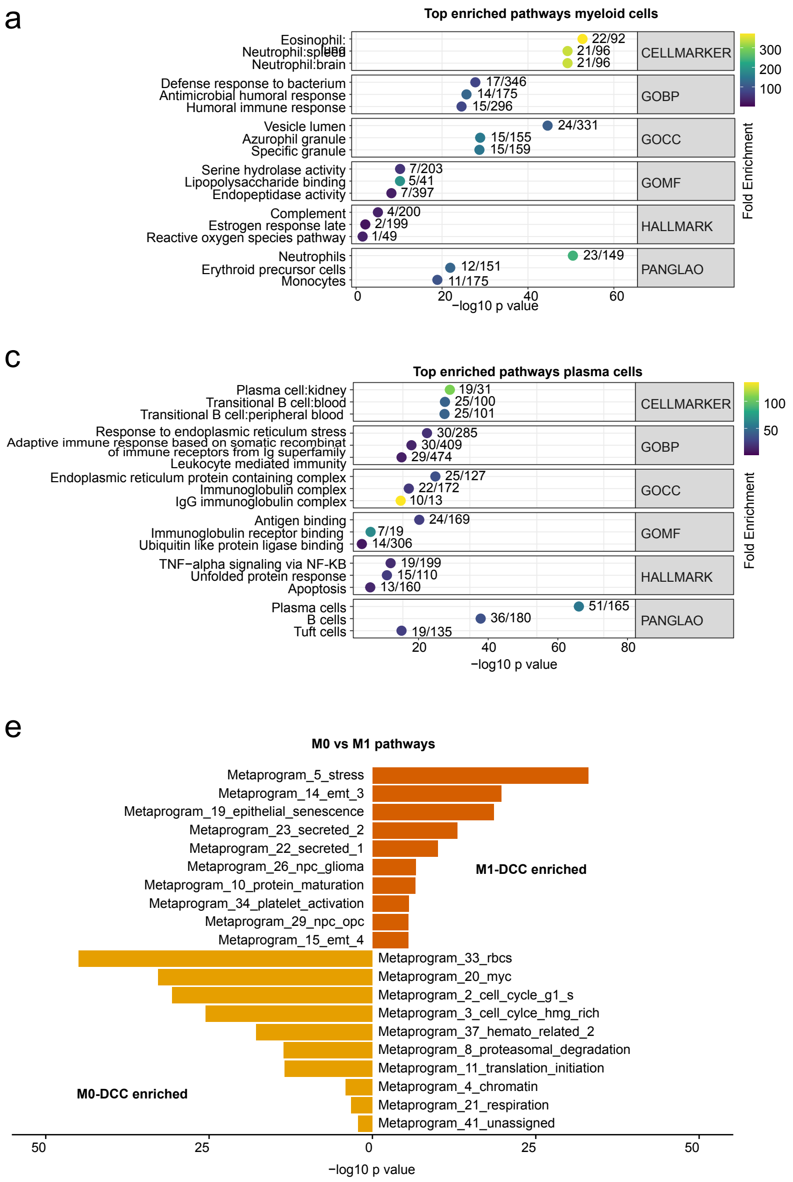

### Supplementary Figure S3

#### Supplementary Figure 4

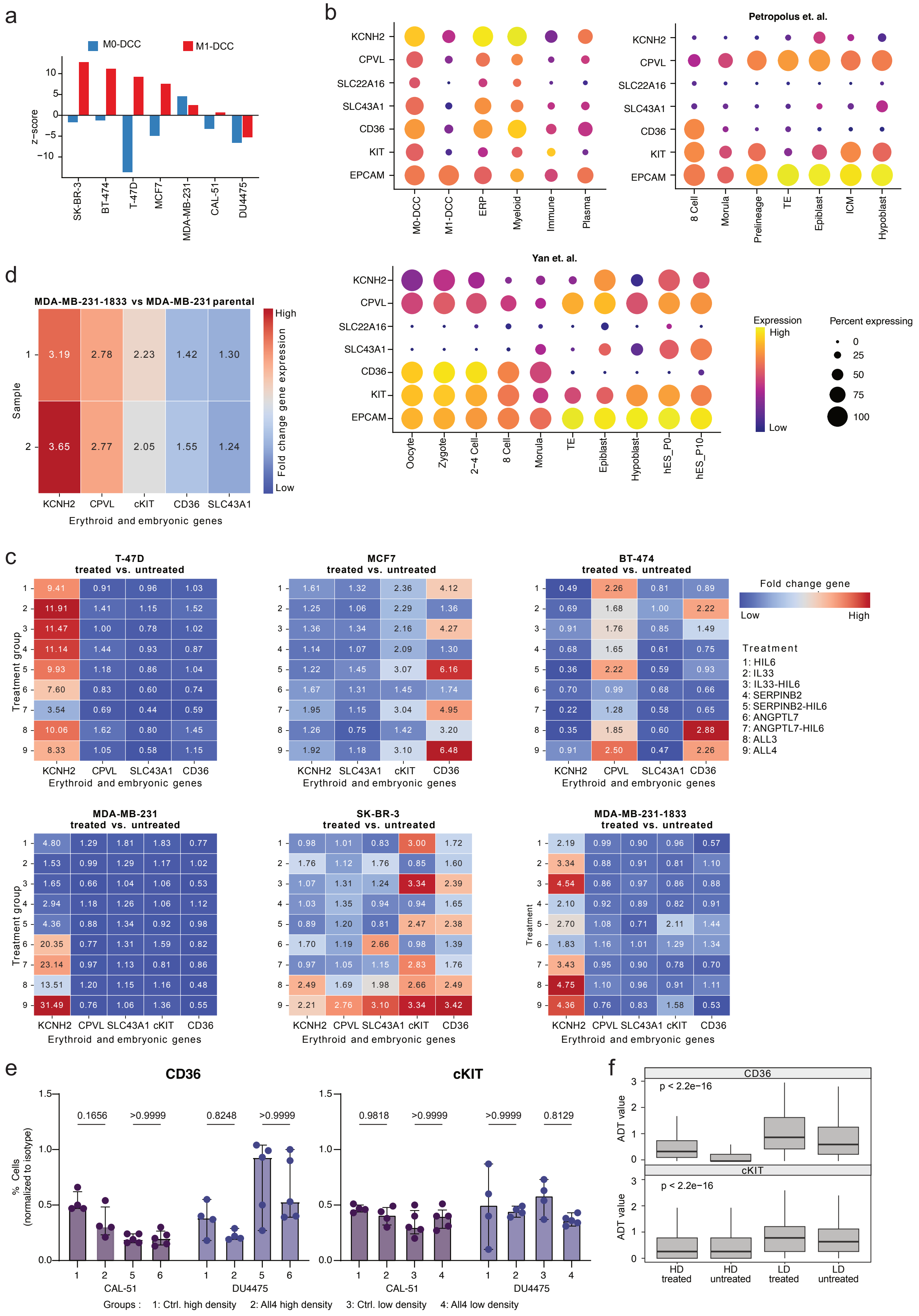

### Supplementary Figure S4

#### Supplementary Figure 3

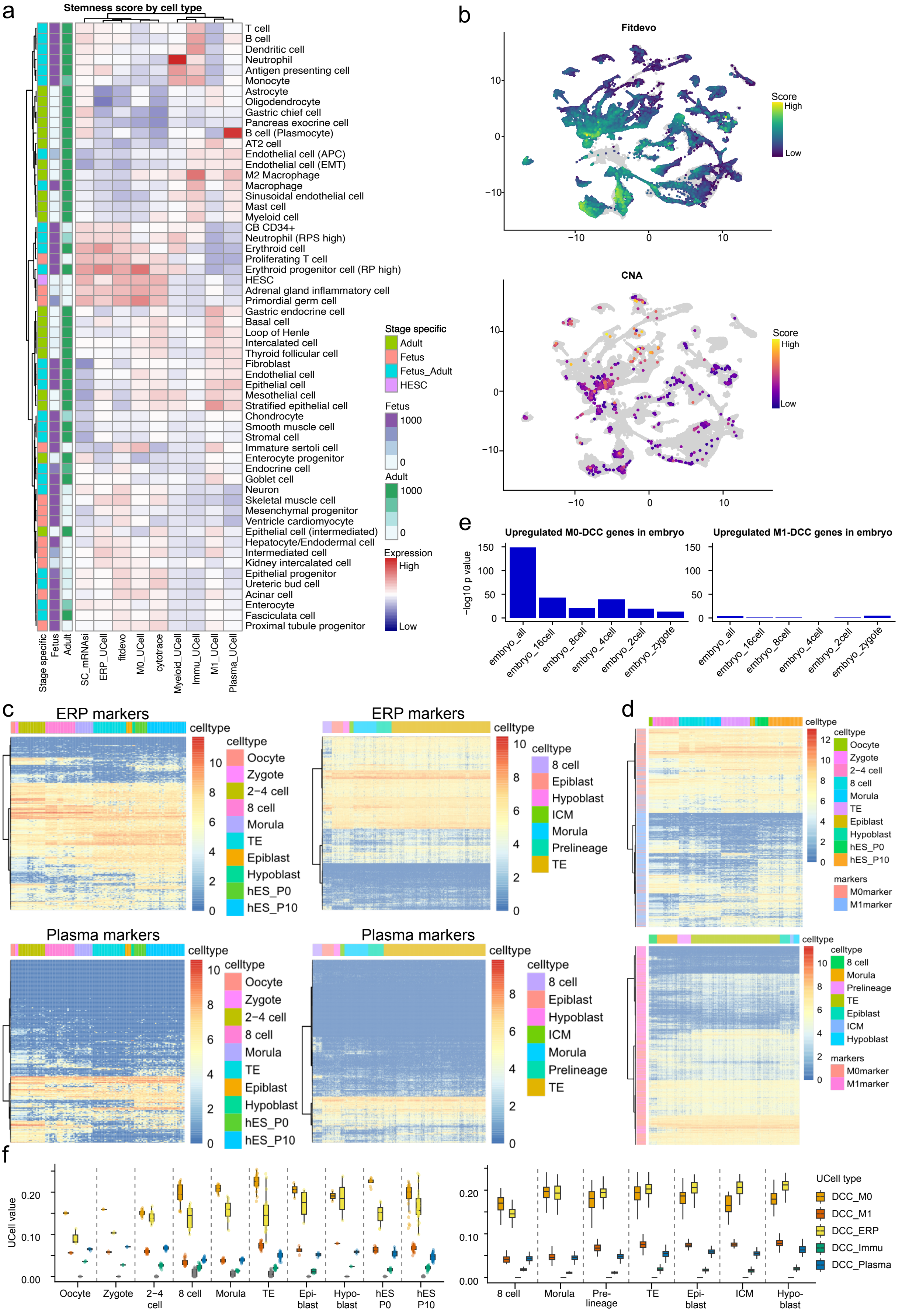
